# Evidence from *Amphioxus* for acquisition of alternative mRNA splicing of NCoR corepressor after its duplication and divergence during vertebrate evolution

**DOI:** 10.1101/2020.03.09.983916

**Authors:** Martin L. Privalsky

**Author notes:** Corresponding author; Address: Department of Microbiology and Molecular Genetics, One Shields Avenue, University of California at Davis, Davis, California 95616.

## Abstract

NCoR-1 and NCoR-2 are transcriptional corepressors encoded in vertebrates by two interrelated loci and play distinct, though overlapping, roles in development, differentiation, and homeostasis. In contrast NCoR is encoded by a single locus in cephalochordates, urochordates, hemichordates, and echinoderms, with vertebrate NCoR-1 and NCoR-2 thought to be the products of a gene duplication originating near the beginning of vertebrate evolution. The structures, molecular properties, and functions of extant NCoR-1 and NCoR-2 are each substantially further diversified by alternative mRNA splicing; however it is unresolved as to whether the alternative-splicing observed in current day vertebrates reflects patterns present in the ancestral common gene or instead arose after the NCoR duplication event. This manuscript reports that *Amphioxus*, a cephalochordate considered representative of the organisms that gave rise to the vertebrate lineage, lacks the alternative NCoR splicing events characteristic of vertebrates. This, together with prior taxonomic comparisons, suggests that the patterns of corepressor splicing found in existing vertebrates arose exclusively after the NCoR duplication event. Further, given that alternative-splicing of NCoR-1 and NCoR-2 appears to have arisen by a mix of convergent and divergent evolution, it is likely that both common and distinct selective pressures were operative on these corepressor paralogs after their divergence.

## INTRODUCTION

NCoR-1 and NCoR-2 are vertebrate coregulator proteins that help mediate repression by a wide variety of transcription factors [1-7]. Vertebrate NCoR-1 and NCoR-2 are encoded by two interrelated loci and play both overlapping and distinct roles in key aspects of development, differentiation, and homeostasis [2,3,8-23]. They share ∼40% amino acid sequence relatedness, a general overall molecular architecture, partially related biochemical properties, and partly overlapping panels of transcription factor partners (Figure 1A, blue vs red schematics). Both NCoR-1 and NCoR-2 operate as bridging proteins that bind to their transcription factor partners through “interaction domains” (IDs) and recruit additional corepressor components through “silencing domains” (SDs), thereby assembling functional corepressor holocomplexes and tethering them to their target genes [1-7]. In contrast to vertebrates only a single NCoR locus has been identified in cephalochordates, urochordates, hemichordates, and echinoderms, with vertebrate NCoR-1 and NCoR-2 thought to represent the products of an ancestral gene duplication and subsequent gene divergence that began near the origin of vertebrate evolution [24].

**Figure 1.**
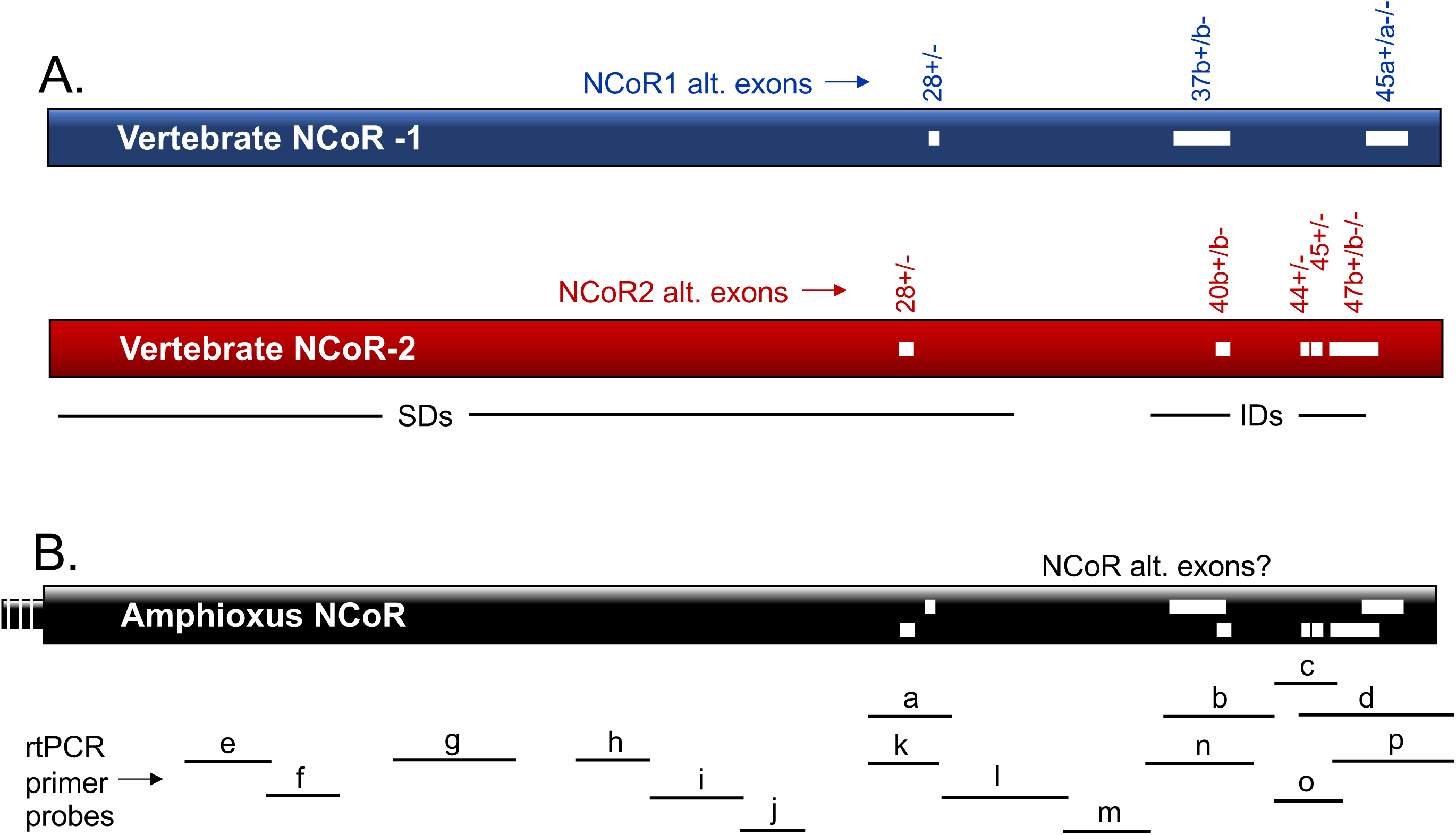
Schematic of vertebrate and Amphioxus NCoR corepressor proteins. Shown are the longest known splice forms of vertebrate NCoR1 and NCoR2 as well as the predicted consensus splice form of *Amphioxus* NCoR. Silencing Domains (SDs) and Interaction Domains (IDs) are depicted. Known alternative spliced exonic sequences are depicted as internal white rectangles within the vertebrate schematics and labeled above by exon number in blue or red. Although also depicted in the *Amphioxus* NCoR schematic for orientation, whether these alternative splice sites are utilized in the cephalochordate is the subject of this investigation. The primer pair probes employed in the RT-PCR experiments are shown below.

Notably both NCoR-1 and NCoR-2 mRNAs are alternatively-spliced in vertebrates to generate a series of distinct corepressor variants (Figure 1A, internal white rectangles) [25-27]; different alternatively-spliced corepressor variants differ in their biochemical properties, engage in different transcription factor partnerships, exhibit different responses to a variety of cellular signaling pathways, and play different, even opposing biological roles [16,28-34]. Comparisons of different taxa suggest that many of these alternative-splicing events arose separately in NCoR-1 and NCoR-2 [35,36]; nonetheless certain of these alternative-splicing events have closely related features that may be indicative of either a retained ancestral splicing event or representative of convergent evolution [24]. To better understand the origins of these events, an examination of alternative-splicing was undertaken in the single NCoR locus found in *Amphioxus* (Figure 1B), a cephalochordate generally thought to be closely related to the ancestral organisms that gave rise to the vertebrate lineage (Figure 2) [37-40]. As reported here, no alternative-splicing was observed in *Amphioxus* NCoR corresponding to the alternative-splicing events found in either vertebrate NCoR-1 or vertebrate NCoR-2. Together with prior taxonomic comparisons the results obtained here indicate that the patterns of corepressor splicing found in existing vertebrates most likely arose separately in NCoR-1 and NCoR-2 after their duplication and divergence and that the apparently related alternative-splicing events found in both paralogs represent convergent evolution.

**Figure 2.**
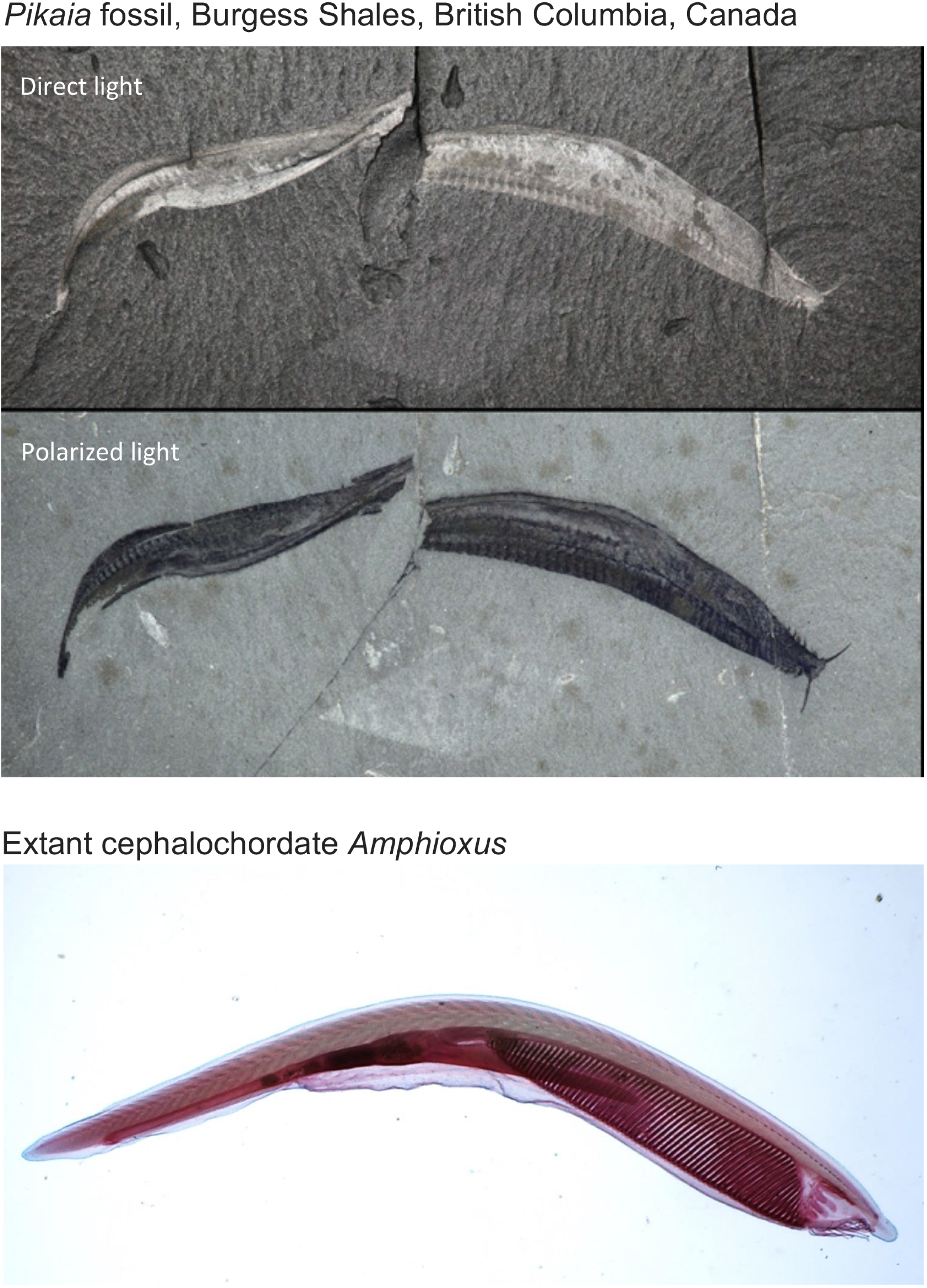
Photographs of a *Pikaia* fossil from the Burgess Shales deposit and a living *Amphioxus* species. Permission was generously provided for use of the fossil images (USNM 83940) by Dr. Thomas Jorstad, National Museum of Natural History USA and for the whole mount image of *Amphioxus* by Dr. Diane Jokinen, Loyola University.

## RESULTS

### Predicted consensus splicing differs between *Amphioxus* and vertebrate NCoRs, yet their overall organization, amino acid sequence, and relevant functional motifs are conserved

The known alternative-splicing sites in vertebrate NCoR-1 and NCoR-2 map within the C-terminal third of their coding region and are particularly prevalent over the ID regions that mediate the interactions of these corepressors with their nuclear receptor transcription factor partners (Figure 1A) [27]. In vertebrate NCoR-1 and N-CoR-2 this region is dispersed over 12 consensus exons but it is condensed into 3 consensus exons in *Amphioxus* NCoR [24]. Nonetheless the overall architecture of the resulting protein product, much of the overall amino acid sequence, and key motifs such as the SD and ID domains are largely conserved between the vertebrate and cephalochordate lineages (Figure 1 and Supplemental Figure S1).

Relatively simple patterns of alternative NCoR-1 and NCoR-2 splicing occur in the more ancient vertebrate taxa, with these patterns conserved and yet more elaborate forms of alternative-splicing added in more recently evolved species [24,35,36]. Overall, three sites of alternative-splicing have been characterized in detail in vertebrate NCoR-1 (exon 28+/28-, exon 37b+/37b-, exon 45a+/45a-/45-) and five in vertebrate NCoR-2 (exon 28+/28-, exon 40b+40/b-, exon 44+/44-, exon 45+/45-, and exon 47b+/47b-/47-) (Figure 1A) [36]. The goal of this work was to determine if *Amphioxus* NCoR exhibited the equivalent of one or more of these alternative-splicing events. Given the possibility of cryptic splice donors and acceptors, alternative-splicing could not be excluded even within the condensed consensus exon structures predicted for the *Amphioxus* NCoR and this region was included in the analysis.

### Alternative mRNA splicing of *Amphioxus* NCoR was not detected using methodology that revealed alternative-splicing in vertebrate N-CoR-1 and NCoR-2

To characterize possible alternative-splicing in *Amphioxus* total RNA was isolated and reverse transcribed into cDNA, with three different tissues/organs studied initially: gonads, hepatic caecum, and muscle. Sequences in the *Amphioxus* NCoR locus corresponding to the alternative-splice sites in vertebrate NCoR-1 and NCoR-2 were identified by multiple-species alignment and used to design suitable primers for use in polymerase chain reaction (Supplemental Figure S1, Table 1). Due to the overlapping nature of NCoR-1 and NCoR-2, four primer pairs, designated a, b, c, and d, were sufficient to test if any of the vertebrate alternative-splicing events also occurred in *Amphioxus* NCoR (Figure 1). The resulting PCR products were analyzed by gel electrophoresis and visualized by ethidium bromide staining. This methodology yields a series of distinct sized PCR products from each alternative-splice site corresponding to each alternatively-spliced mRNA produced at that site. This method is both sensitive and reproducible, as detailed in Materials and Methods.

A strong PCR product was detected using *Amphioxus* cDNA from each of the three tissues and employing primer pairs flanking each of the alternative-splice sites characterized in vertebrate NCoR-1 or NCoR-2 (Figure 3) with only one exception: hepatic caecum cDNA and primer pair d yielded only a very faint PCR product (Figure 3). Notably the appropriately sized PCR product could be observed on extended exposure (Figure 3, insert). Probing the hepatic caecum cDNA with additional primer pairs further confirmed that this tissue expresses the anticipated NCoR mRNA sequences and no detectable alternatively-spliced variants over this region (Figure 5 and data not shown).

**Figure 3.**
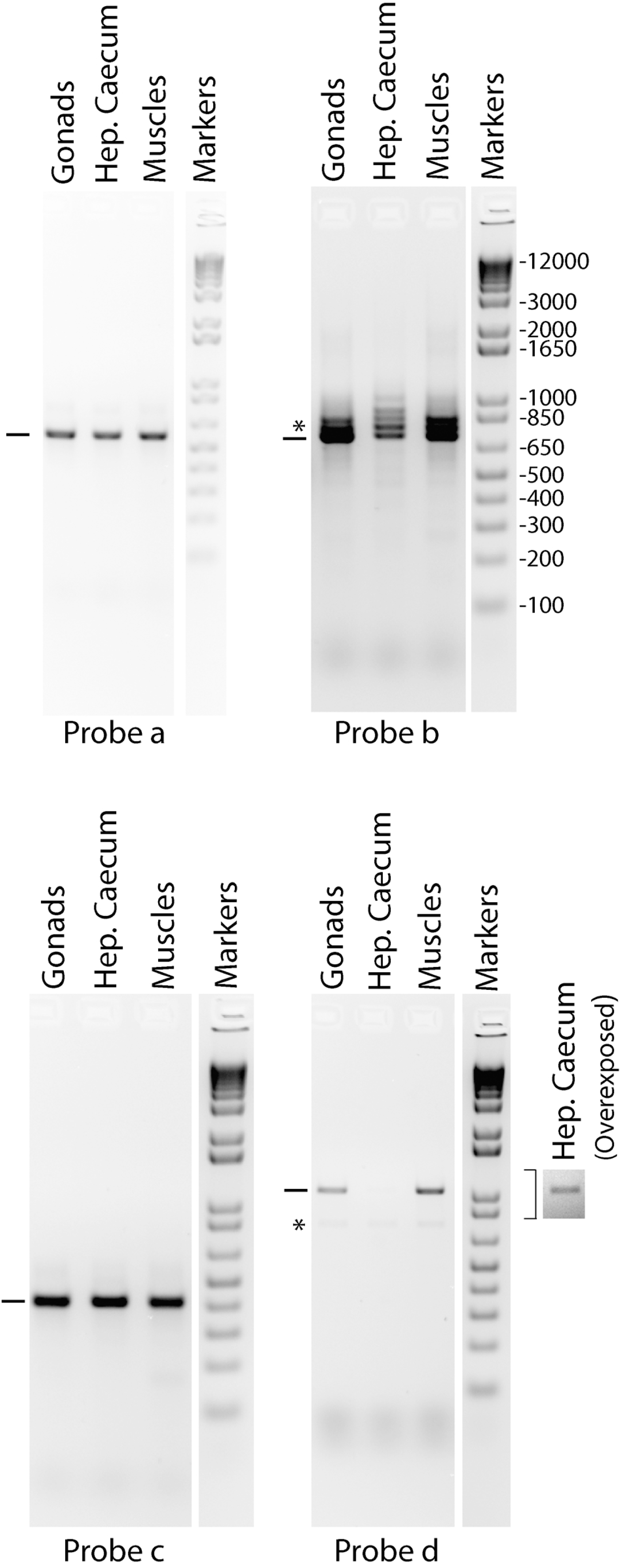
RT-PCR analysis of mRNAs from different *B. floridae* tissues; regions of NCoR locus alternatively-spliced in vertebrate NCoR1 or NCoR2. mRNAs from the three tissues indicated were tested. The locations of the primer pair probes are indicated in Figure 1. The PCR products from probes b and c and the associated molecular weight markers were resolved on a single gel; this one marker lane was therefore duplicated and attached to both of these panels. Inset: over-exposed and enhanced probe d RT-PCR from hepatic caecum. Asterisks indicate background bands generated by misprimings on reiterative DNA sequences upstream of the probe b hybridization site or amplification by primer pair d of an *Amphioxus* gene transcript unrelated to NCoR.

**Figure 4.**
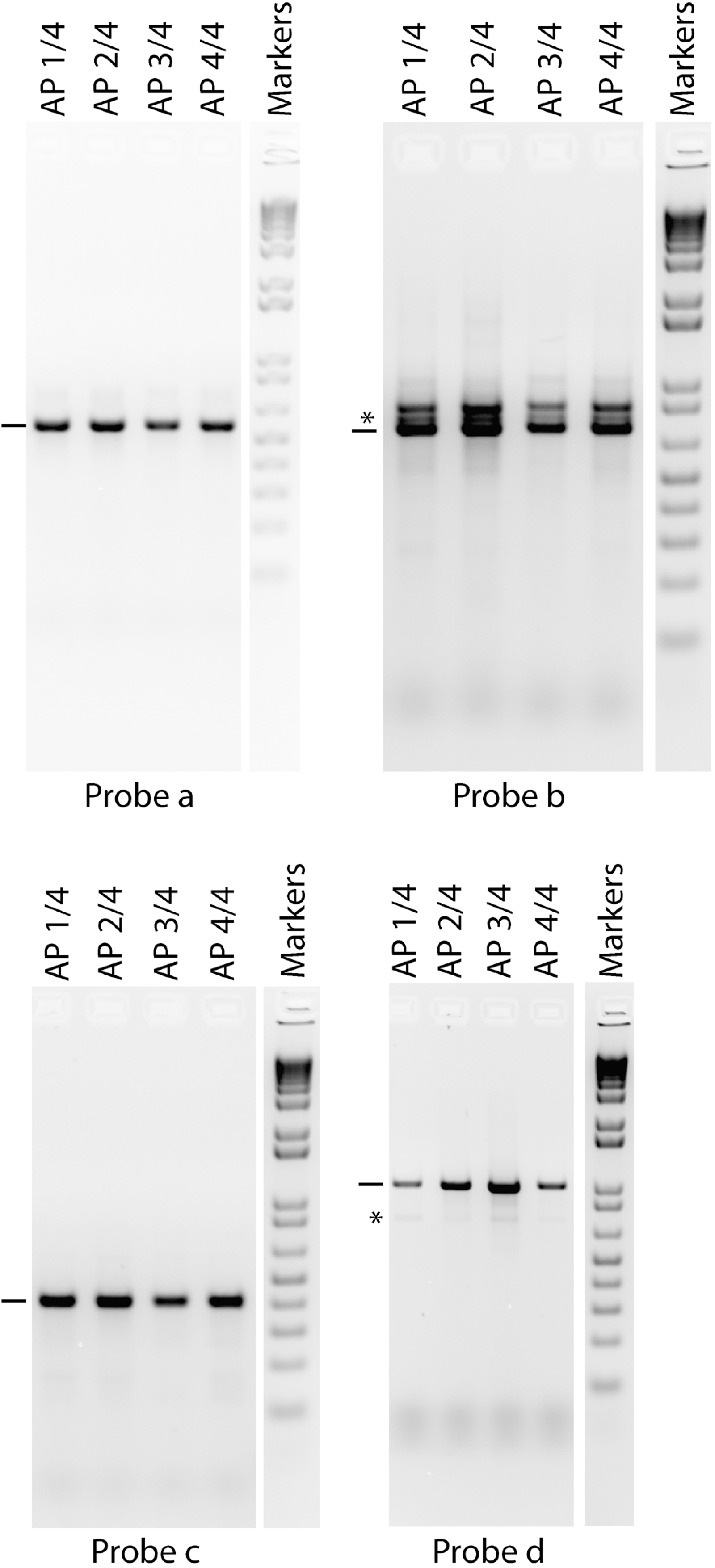
RT-PCR analysis of mRNAs from different *B. floridae* body sections; regions of NCoR locus alternatively-spliced in vertebrate NCoR1 or NCoR2. mRNAs from B. *floridae* body sections AP1. AP2, AP3, and AP4 were tested. The locations of the primer pair probes are indicated in Figure 1. The PCR products from probes b and c and the associated molecular weight markers were resolved on a single gel; this one marker lane was therefore duplicated and attached to both of these panels. These samples and molecular weight markers were also resolved on the same gel as in Figure 3; the molecular weight marker lane for probes b and c is therefore also identical in both figures. Asterisks indicate background bands generated by misprimings on reiterative DNA sequences upstream of the probe b hybridization site or amplification by primer pair d of an *Amphioxus* gene transcript unrelated to NCoR.

**Figure 5.**
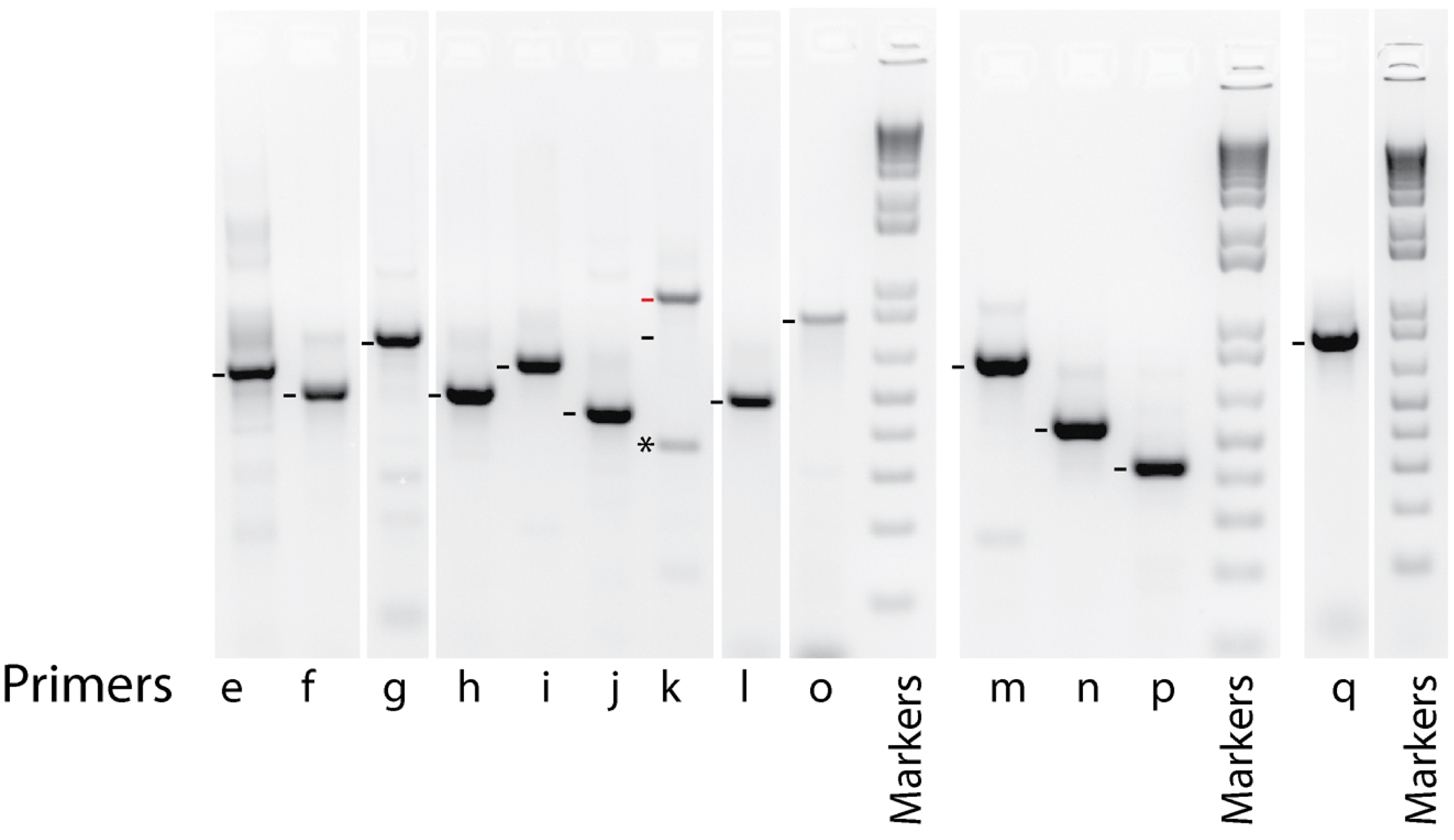
RT-PCR analysis of mRNAs from different *B. floridae* sections; general survey of NCoR transcript. mRNAs from pooled B. *floridae* body sections AP1, AP2, AP3, and AP4 were tested. The locations of the primer pair probes are indicated in Figure 1. Lanes 1 to 9 shared the molecular weight markers in lane 10; lanes 11 to 13 shared the molecular weight markers in lane 14; lane 15 utilized the molecular weight marker in lane 16. Red bar in lane 7 indicates actual 943bp RT-PCR product (red bar); black bar indicates 684bp predicted by the consensus splicing (see text). Asterisk indicates a background band generated from *Amphioxus* gene transcripts unrelated to NCoR.

In each case the PCR products corresponded in size to the full-length mRNAs predicted from these sites with no products attributable to alternative-splicing detected in any of the primer pair/cDNA reactions (Figure 3). The identities of the PCR products were confirmed by DNA sequence analysis and/or use of additional primers (data not shown). Several minor PCR products were observed in certain cDNA/primer reactions (*) but were determined to be artifacts by the same means; these typically represented misprimings within reiterative DNA sequences or primings on partially related sequences mapping to non-NCoR sequences elsewhere in the *Amphioxus* genome (see Figure legends). No PCR products were detected if reverse transcriptase was omitted from the cDNA synthesis reaction (data not shown), confirming the absence of genomic DNA contamination.

### No vertebrate-like alternative-splicing was detected in NCoR in a broader survey of *Amphioxus* tissues

To test if alternative-splicing might occur in tissues others than those tested above, *Amphioxus* were dissected into 4 broad anterior to posterior segments (denoted AP1/4, AP2/4, AP3/4, and AP4/4); mRNA was isolated from each section and was subjected to the same RT-PCR analysis as above (Figure 4). All 4 sections yielded results identical to the individual tissues studied above: single RT-PCR products corresponding to the full-length transcripts predicted, with no evidence of alternative-splicing detected over these regions (Figure 4).

### A wider survey using ordered sets of primer pairs did not detect alternative mRNA splicing in other regions of Amphioxus NcoR

The RT-PCR methodology was repeated using cDNAs pooled from the AP1/4, AP2/4, AP3/4, and AP4/4 sections and a series of ordered primer pairs designed to probe the *Amphioxus* transcript more broadly (Figure 1B). Only RT-PCR products corresponding to the full-length consensus transcript were detected with a single exception (Figure 5). Probe k amplified a 943bp product (red bar) instead of the 684bp (black bar) product predicted by the consensus splicing. Sequence analysis revealed inclusion of an additional 185bp and 74 bp of genomic sequence at consensus splice boundaries 5270bp and 5478bp within the predicted *Amphioxus* transcript (data not shown). Notably none of the predicted consensus 684bp product, nor any additional NCoR-related product, was observed indicating that this phenomenon was not alternative-splicing resulting in multiple transcripts but rather non-consensus splicing resulting in a single transcript. These results indicate that the cephalochordate NCoR mRNA is not detectably alternatively-spliced within the limits of this survey (please see the Discussion).

## DISCUSSION

The duplication and divergence of NCoR into NCoR-1 and NCoR-2 during vertebrate evolution enhanced the diversity of actions available to these paralogs verses the ancestral single copy gene, with NCoR-1 and NCoR-2 displaying distinct molecular properties and mediating distinct biological roles [1-7]. Further, vertebrate NCoR-1 and NCoR-2 are both alternatively-spliced to create an extensive series of corepressor protein variants that differ in abundance in different tissues, preferentially associate with different transcription factor partners, respond to signal transduction pathways in divergent ways, and play distinct, even opposing, biological roles [16,28-34]. Thus alternative-splicing further broadens the diversity created by the duplication and divergence of the NCoR locus itself. Understanding the appearance of these alternative-splicing events over evolution therefore provides insights into the biological roles of the different extant splice variants and how they responded to different selective pressures in different species.

The sites and patterns of species-specific expression of six of the vertebrate alternative-splice sites are consistent with their arising independently by divergent evolution in the two paralogs: NCoR-1 exon 28+/28-, NCoR-1 exon 45a+/45a-/45-, NCoR-2 28b+/28b- (unrelated to NCoR-1 exon 28b+/b-in location and sequence), NCoR-2 exon 44+/44-, NCoR-2 45+/45-, and NCoR-2 exon 47b+/47b-/47-. In contrast the NCoR-1 exon 37b+/37b- and NCoR-2 exon 40b+/40b-alternative-splicing events map within overlapping, interrelated sequences within these two paralogs and insert or omit a conserved “ID3” domain in both that controls their interaction with their nuclear receptor/transcription factor partners. The NCoR-1 exon 37b+/37b-alternative-splice is observed exclusively in placental mammals whereas the NCoR-2 exon 40b+/40b-is present in all vertebrates examined [35,36]. Although this is suggestive of a later evolutionary acquisition of the former, and thus convergent evolution, it was also possible that these alternative-splice events preceded the NCoR duplication event but were selectively lost in NCoR-1 until their reappearance later in evolution [24]. It was therefore decided to further evaluate this phenomenon and the possibility of other examples of vertebrate-like alternative-splicing in *Amphioxus*.

The experiments reported here demonstrated that neither the NCoR-1 exon 37b+/37b-nor the NCoR-2 exon 40b+/40b-variants, nor any of the other vertebrate alternative-splicing events studied here, were detected in *Amphioxus*. Taken together with the taxonomic analyses previously reported these results provide additional support for the proposal that the alternative-splicing observed in extant NCoR-1 and NCoR-2 arose independently in these corepressor paralogs after their duplication and subsequent divergence during vertebrate evolution. Given both divergent and convergent events appear to have occurred in these different paralogs, this conclusion further suggests that NCoR-1 and NCoR-2 have been subject to a mix of shared and distinct selective pressures that differed in the different vertebrate lineages. Notably in this regard NCoR 37b+/37b-alternative splicing plays a key role in murine lipid and glucose metabolism [32,34]; it is tempting to speculate that its convergent acquisition in placental mammals after the acquisition of the NCoR-2 exon 40b+/40b-alternative splice reflects the unique metabolic needs of eutherian mammals.

A more extensive series of primer pairs surveying broader regions of *Amphioxus* mRNA failed to detect alternative splicing even outside of the regions that correspond to alternative splice sites in NCoR-1 or NCoR-2 in vertebrates. However the current survey was not comprehensive and it remains possible that alternative splicing events exist in *Amphioxus* NCoR that are unique to this cephalochordate and that do not correspond to those in vertebrates.

Limitations to these studies: although our methodology can detect alternatively-spliced products as low as? 5% of total it cannot be excluded that some splice variants may be expressed at levels below this limit; such low levels of expression, if they occur, would be of unclear biological significance. Similarly although no alternative mRNA splicing was detected in any of the tissues tested, including broad sections of the *Amphioxus* body plan, it is conceptually possible that *Amphioxus* NCoR mRNA is alternatively-spliced in a very minor tissue or cell type such that its expression level is diluted within the broader tissues samples to below the detection limit. The hypothetical inclusion of very large introns into the *Amphioxus* consensus splicing pattern (i.e. significantly longer than those seen in vertebrates) might be difficult to detect given the reduced efficiency of Taq polymerase on very long templates. Finally given that a single mutation can eliminate a splice donor or acceptor site [24] it remains possible that the *Amphioxus* lineage lost one or more alternative splicing events during its evolution that were operative in the pre-vertebrate predecessor. Nonetheless the simplest interpretation of the data as a whole supports the proposal that the alternative slicing observed in extant NCoR-1 and NCoR-2 arose only after the appearance of vertebrates.

## MATERIALS AND METHODS

### Sources, isolation of RNA, cDNA synthesis, and PCR analysis

Adult *Amphioxus* (*Branchiostoma fluoridae*) were obtained from Gulf Specimen Marine Laboratories. *Amphioxus* were euthanized by immersion in 200 mg/liter Triciane methanesulfonate (MS-22) in sea water until all movement ceased for a minimum of 10 minutes, followed by decapitation and mechanical ablation of the neural bulge. The animals were then dissected into four sections from anterior to posterior, or the gametes, hepatic caecum, and muscles were individually removed. Samples were stored at −80 °C in RNAlater (Life Technologies, Grand Island, NY). Total RNA was subsequently isolated from approximately 30 μg of each tissue using “bead beating” homogenization and an RNAeasy mini-kit (Qiagen, Hilden, Germany) following manufacturer’s directions.

Up to 1 μg of RNA from each sample was converted into cDNA using a QuantiTect Reverse Transcription Kit and manufacturer’s directions (Qiagen, Hilden, Germany); a DNA-wipeout pre-step was included to avoid genomic contamination. Two to four μl of each 120 μl cDNA preparation were amplified by PCR using oligonucleotide primer pairs flanking each alternative splice-site (Supplemental Table S1), GoTaq enzyme, GoTaq buffer, and the manufacturer’s recommendations (Promega, Madison WI). PCR was typically performed at 94 °C for 2 mins. followed by 34 cycles of 30-60 secs. at 94 °C, 30-60 secs. at 53 °C, 60-120 secs. at 72 °C, followed by a 5 min. final extension step at 72 °C. The PCR products from each primer pair were resolved by electrophoresis in a 2% agarose gel in 1X Tris-Acetate-Ethylenediaminetetraacetate buffer, the DNA bands were visualized with ethidium bromide, and were documented by use of a digital camera. NCoR-2 was previously referred to as SMRT (an alternative nomenclature); the exon numbering system here is as in [35].

### Accuracy of the RT-PCR methodology

Although not favored for quantitation of absolute levels of nucleic acids, the end-point RT-PCR technique as adapted here is reproducible, sensitive, and accurate for the determination of the relative abundances of the different splice variants produced at a given alternative-splice site. A more detailed justification is presented in [36], but briefly: (a) The RT-PCR methodology is internally-controlled, with all splice variants from a given splice site amplified in parallel in the same tube under identical conditions using the same primers and analyzed in a single lane in an electrophoresis gel. This method is relatively insensitive to non-uniformities in primer or reaction efficiency, pipetting, temperature, ramp-rate, lane-loading, electrophoresis, staining or destaining. (b) The primers were designed to minimize as much as possible hybridization temperature and size differences between the potential alternative-splice products produced from each splice site (with the goal of minimizing differences in the efficiency of their amplification). (c) Experimental tests of this approach yielded high reproducibility and low standard deviations in repeated experiments. For mRNAs that are alternatively-spliced the relative ratios of the corresponding RT-PCR products from each given alternatively-spliced site are not significantly altered over a wide range of cycle numbers and input cDNA concentrations.

## DECLARATIONS

### Ethical approvals and considerations

Although *Amphioxus* do not fall under the guidelines required for use of vertebrate animals all studies reported here were carried out under the vertebrate animal guidelines and restrictions mandated by the University of California at Davis Animal Use committee for teleost fish. This work has not been submitted or published elsewhere.

### Author’s contributions

MLP performed the experiments, the analysis, and the assembly and writing of this manuscript.

## Acknowledgements

The author is sincerely grateful to Dr. Michael L. Goodson for having originally pioneered the RT-PCR methodology and for helpful discussions, and to Drs. Thomas Jorstad and Diane Jokinen for permission to use the images in Figure 2.

## SUPPLEMENTAL MATERIALS

**Supplemental Table S1.**
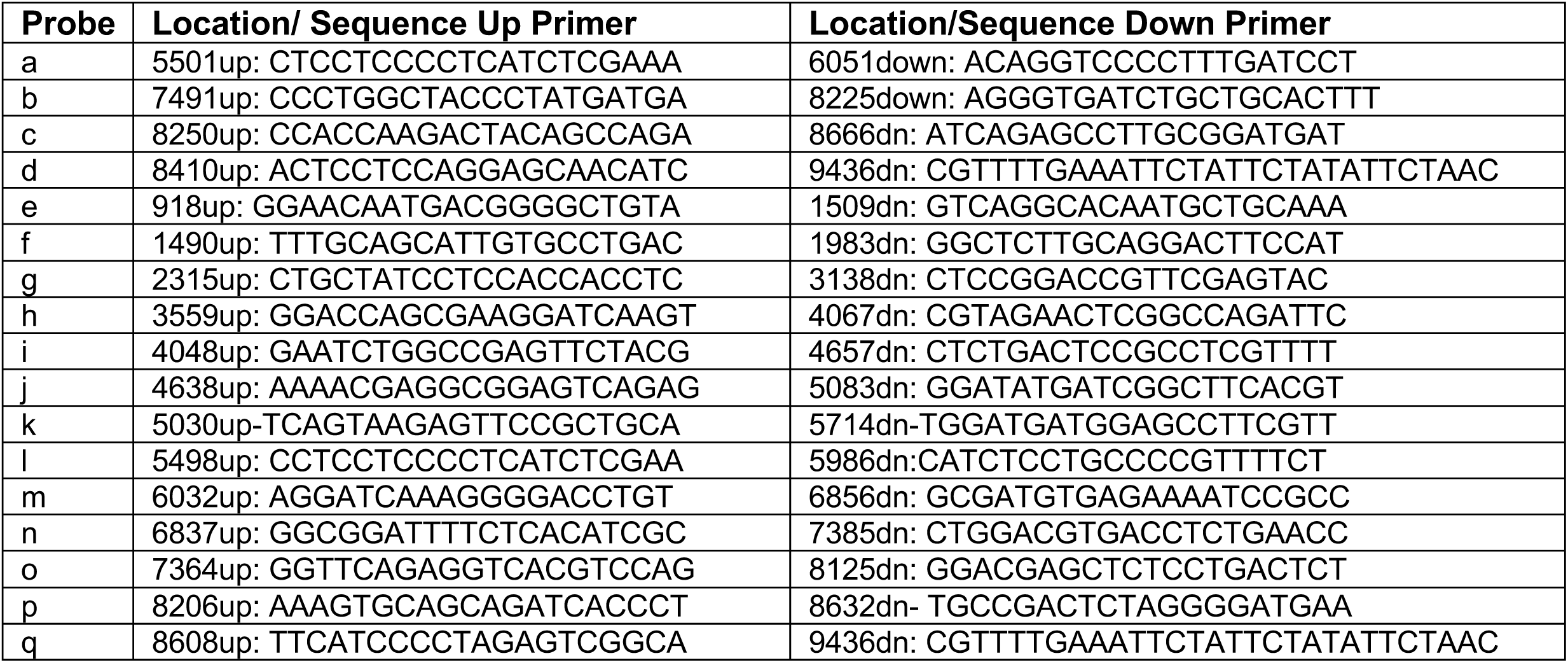
Primers employed in RT-PCR. Oligonucleotide sequence and position in *Amphioxus B. floridae* cDNA.

**Supplemental Figure 1.**
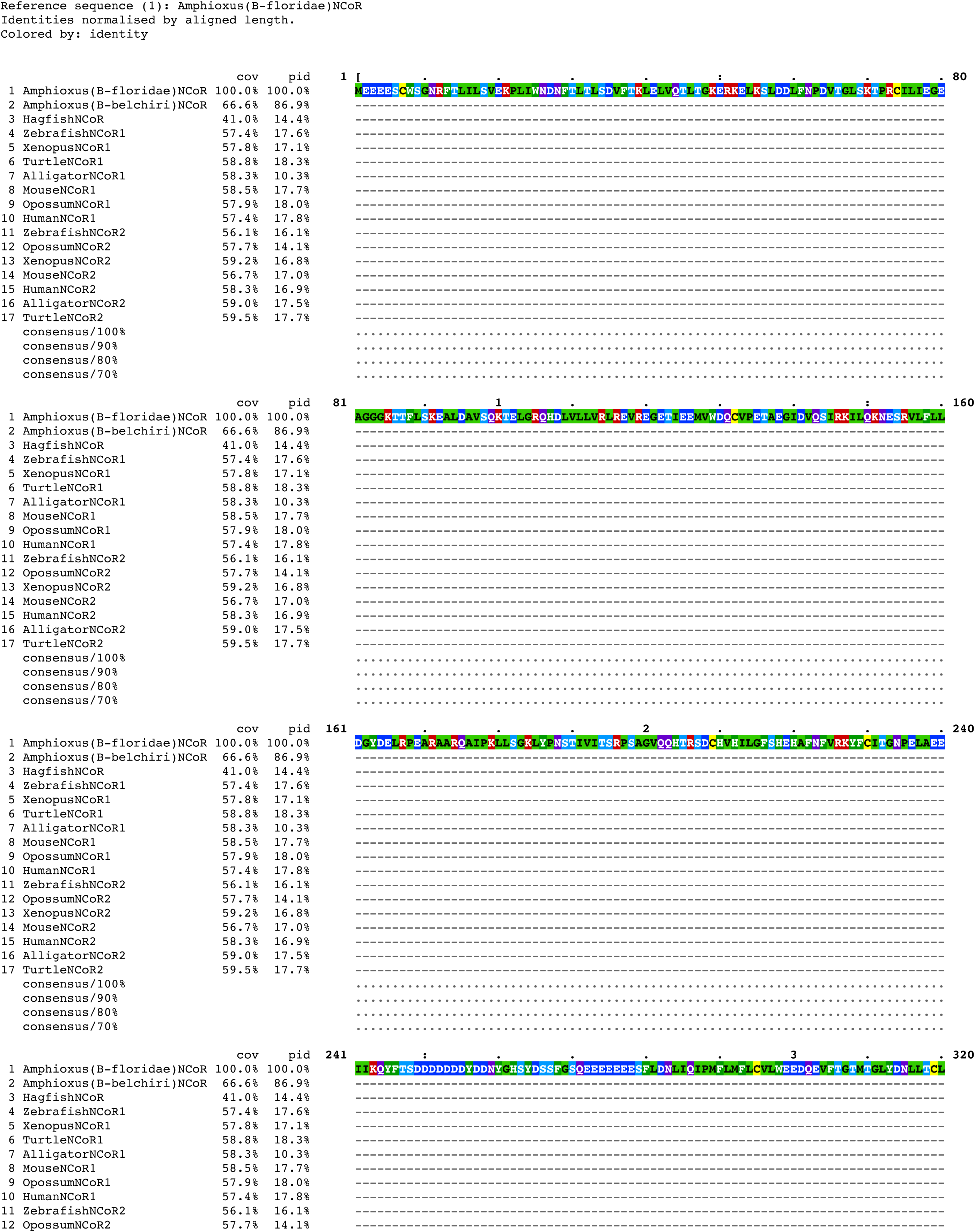
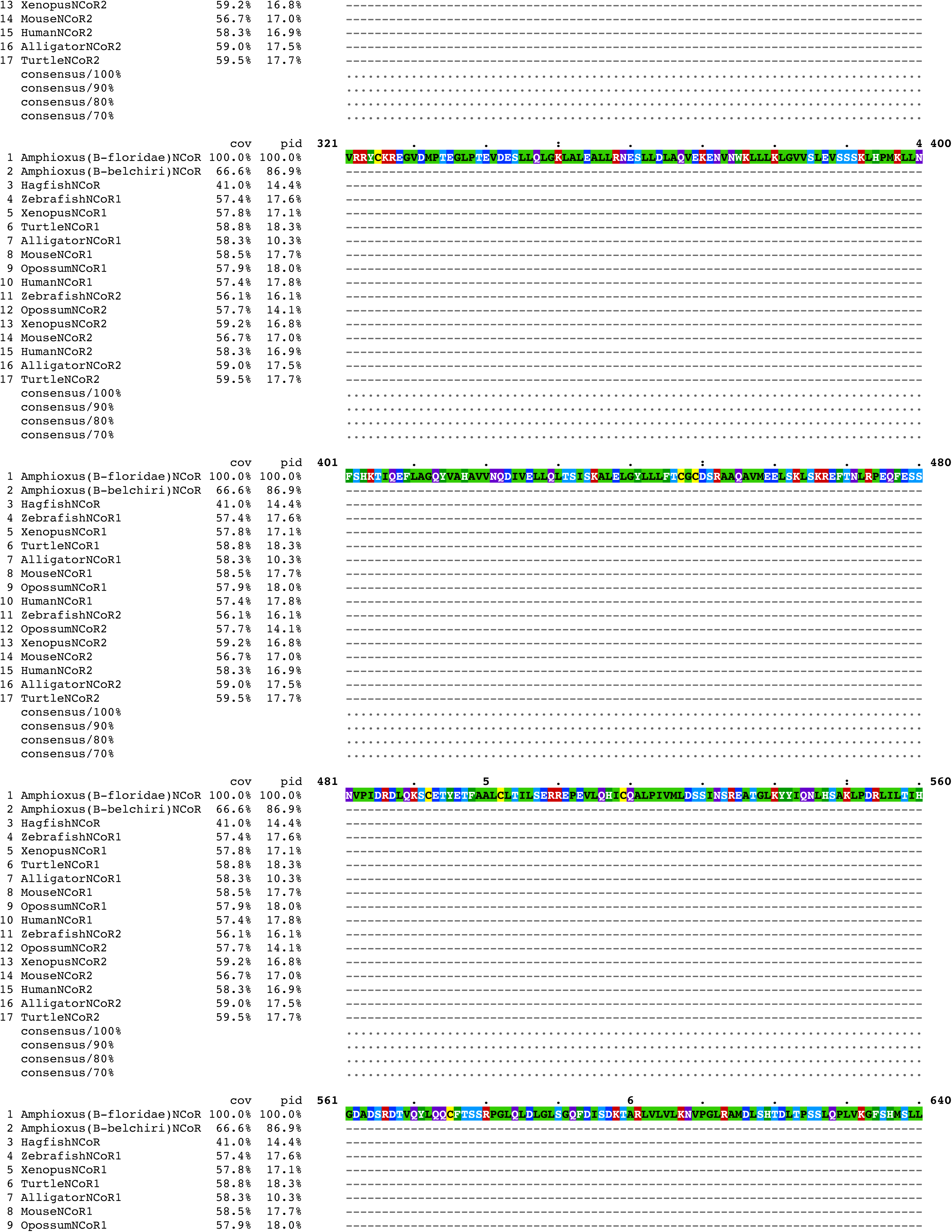
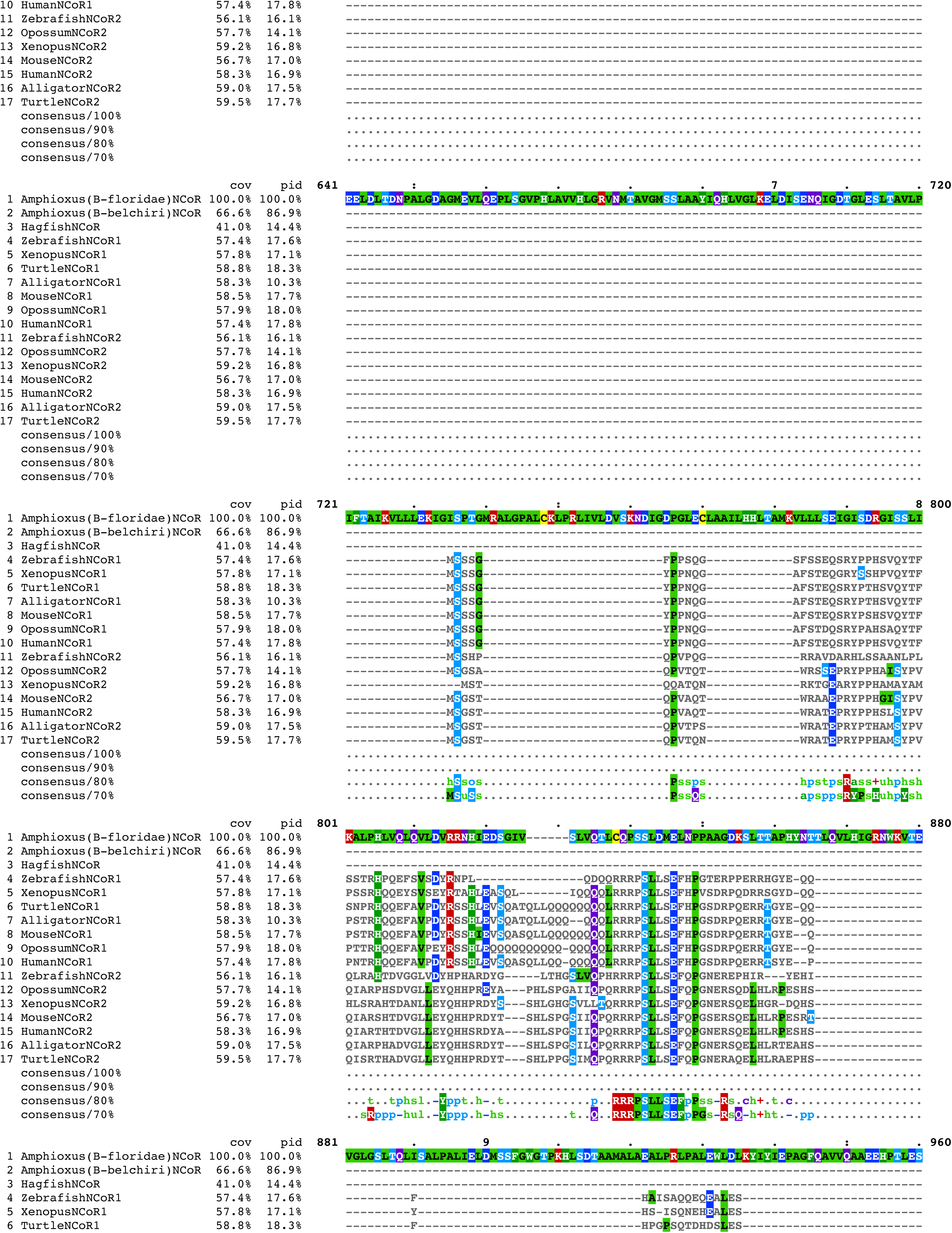
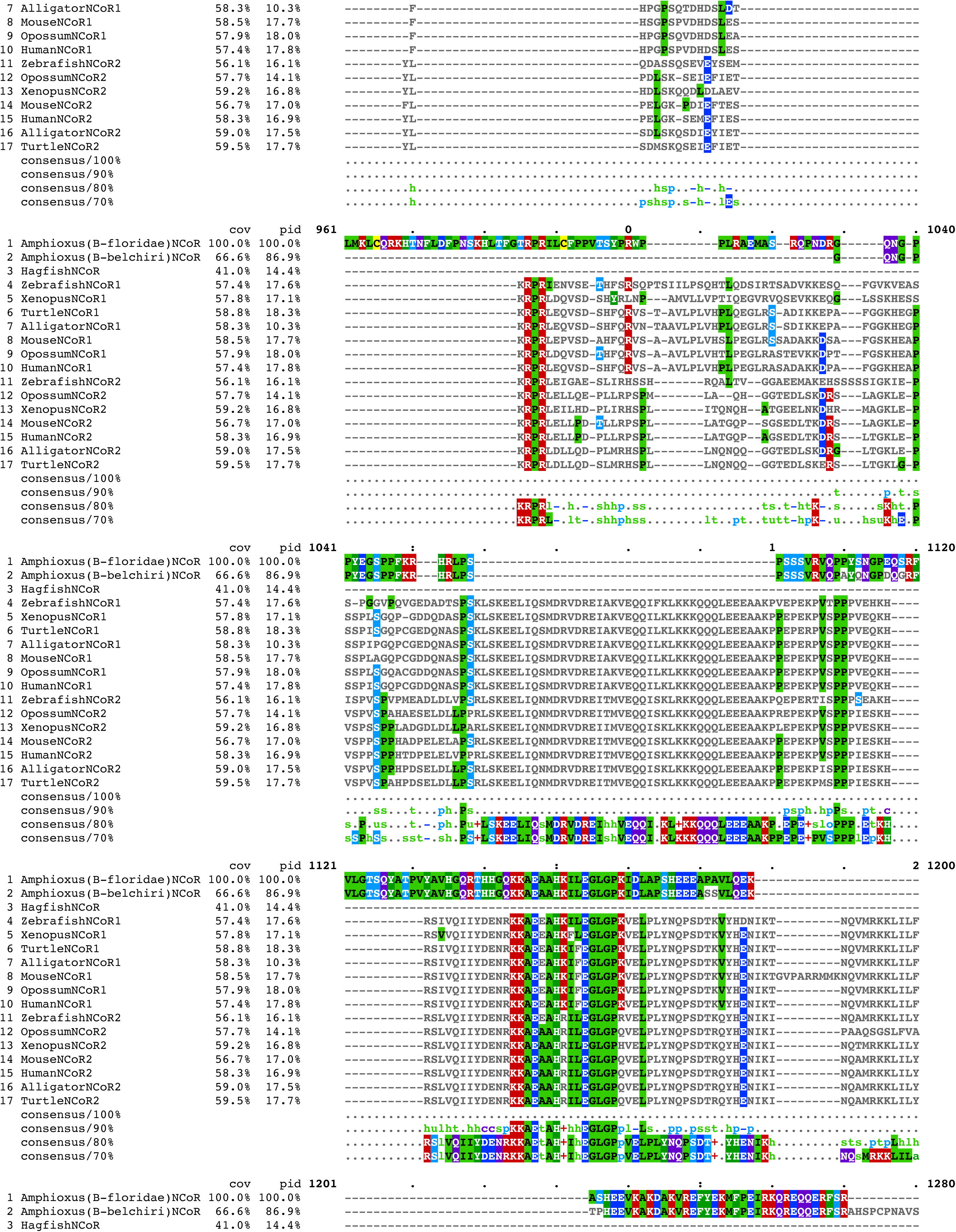
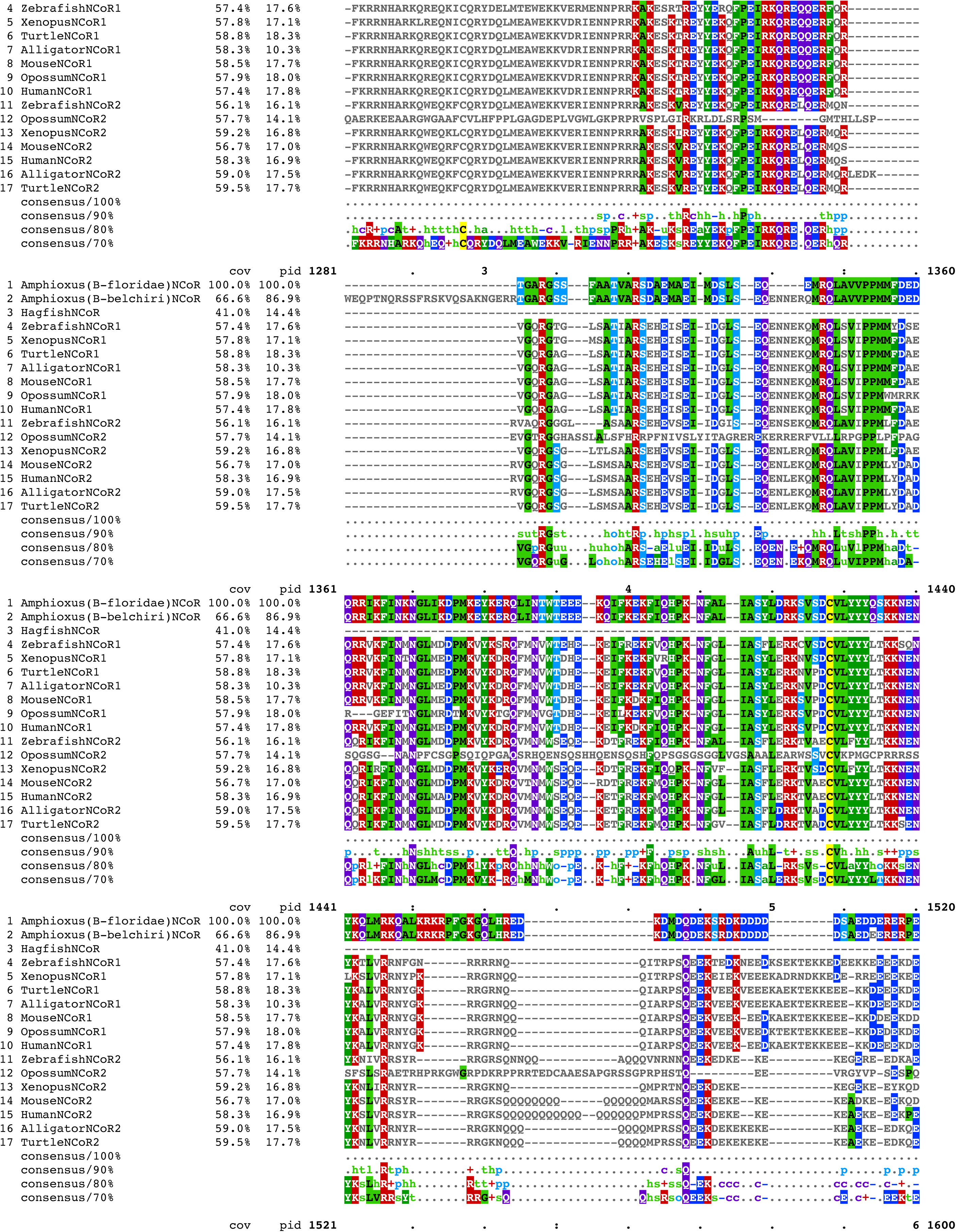
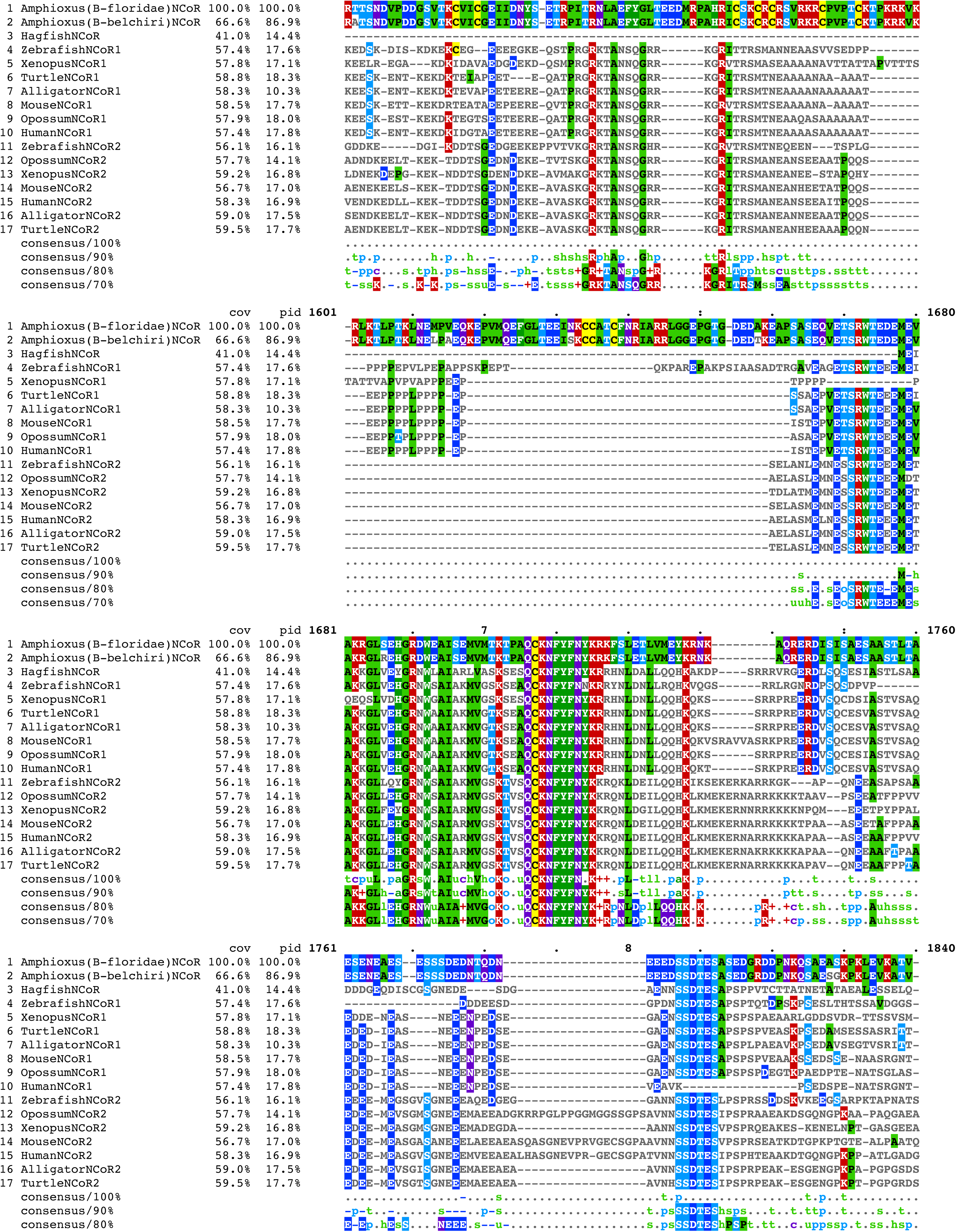
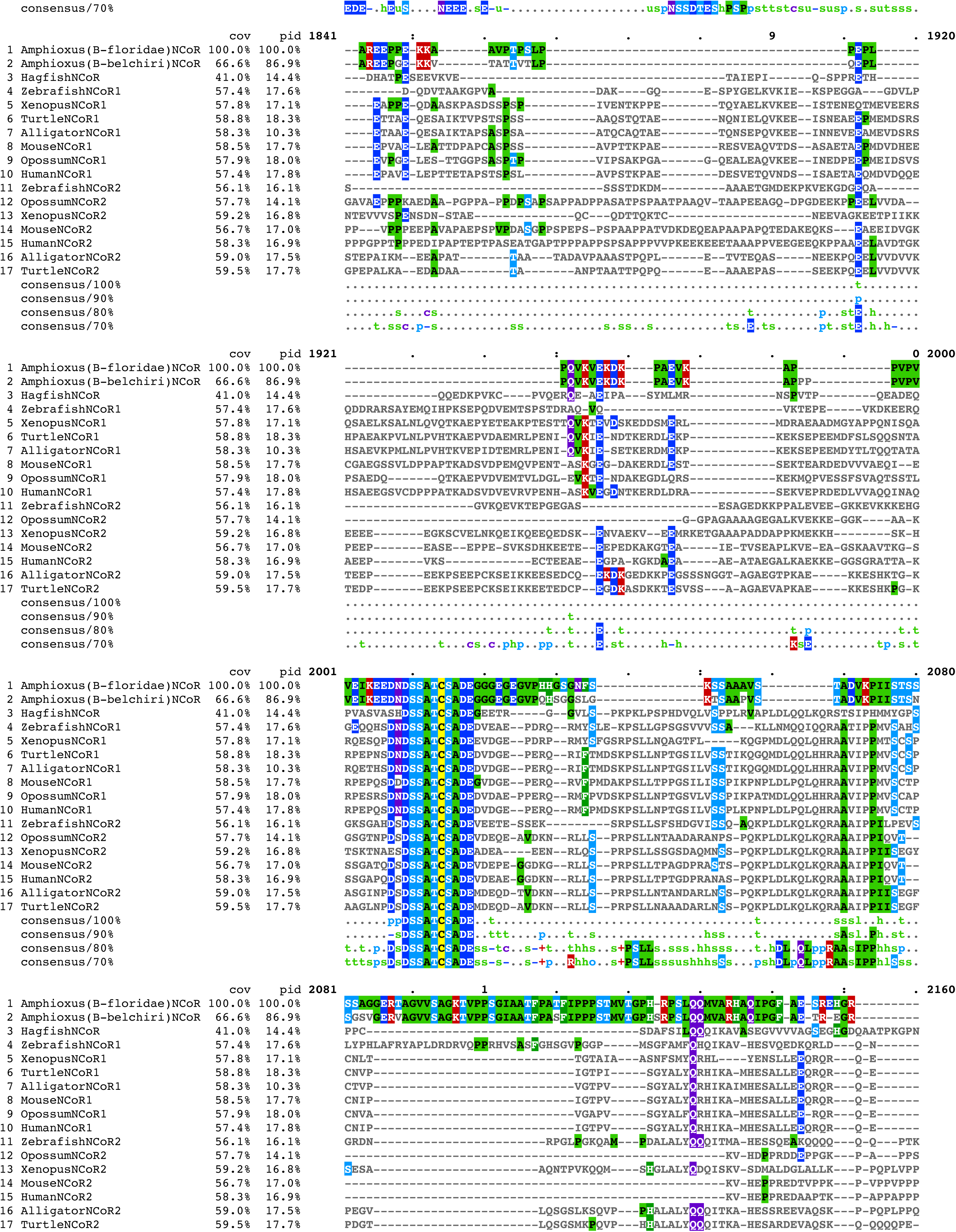
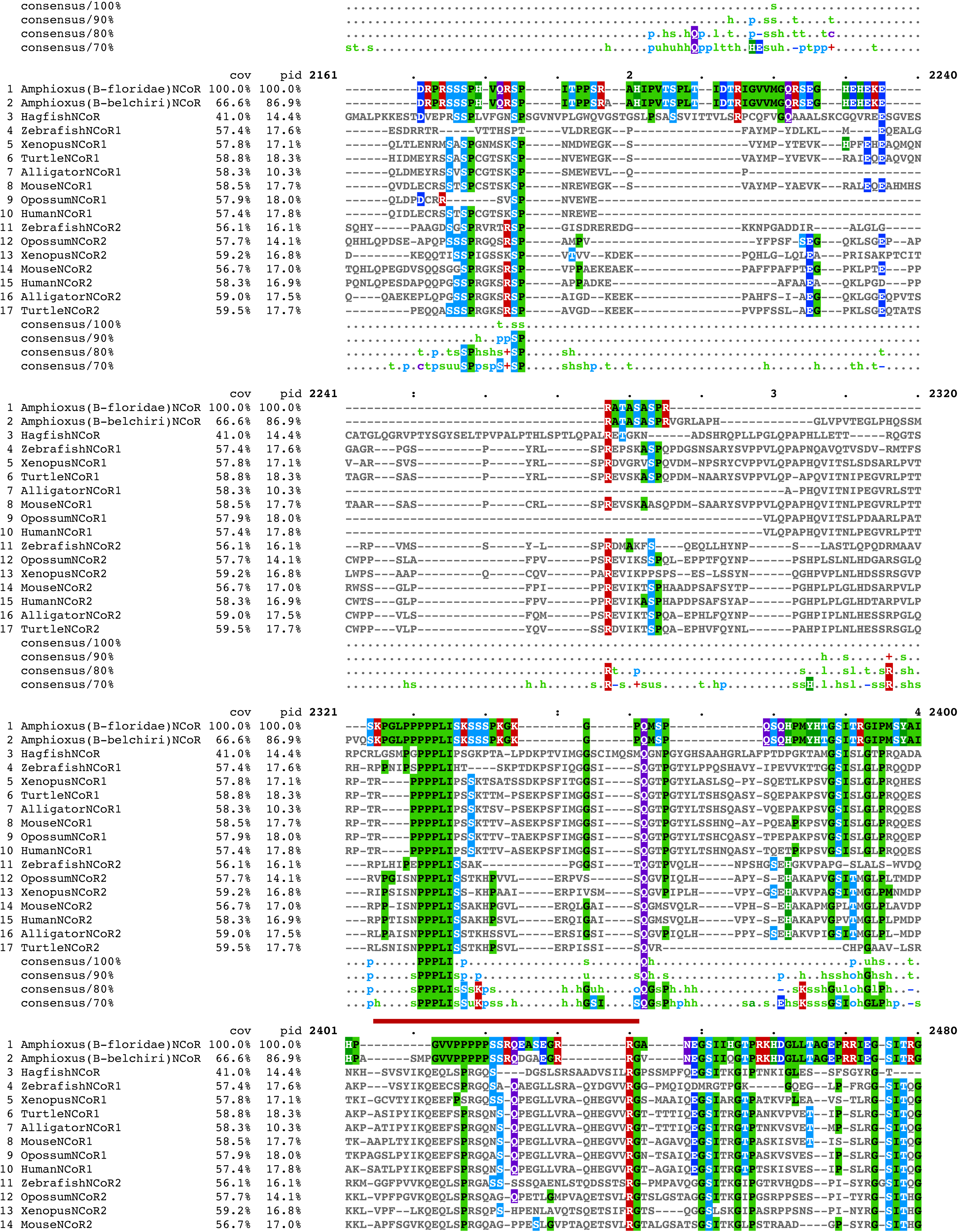
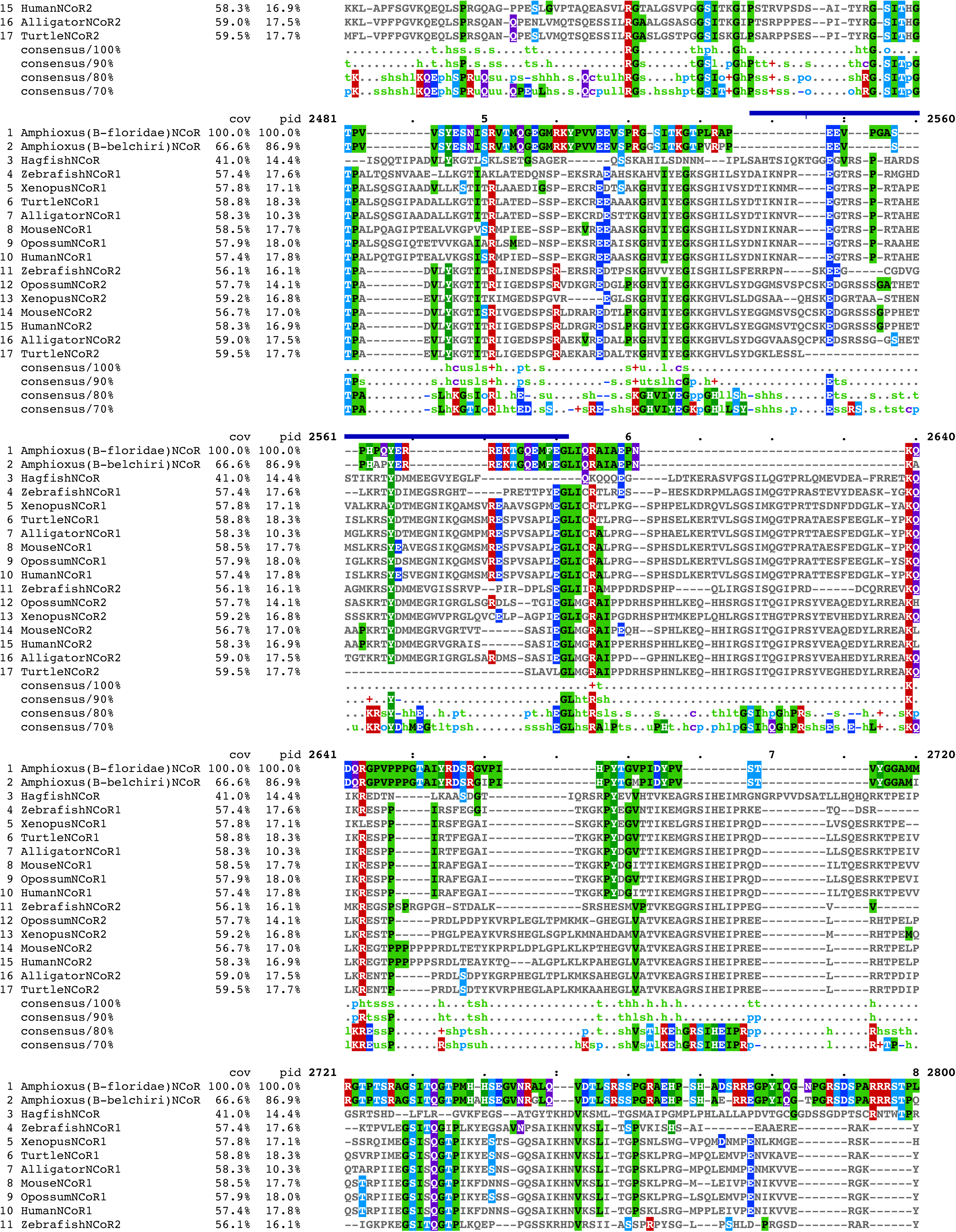
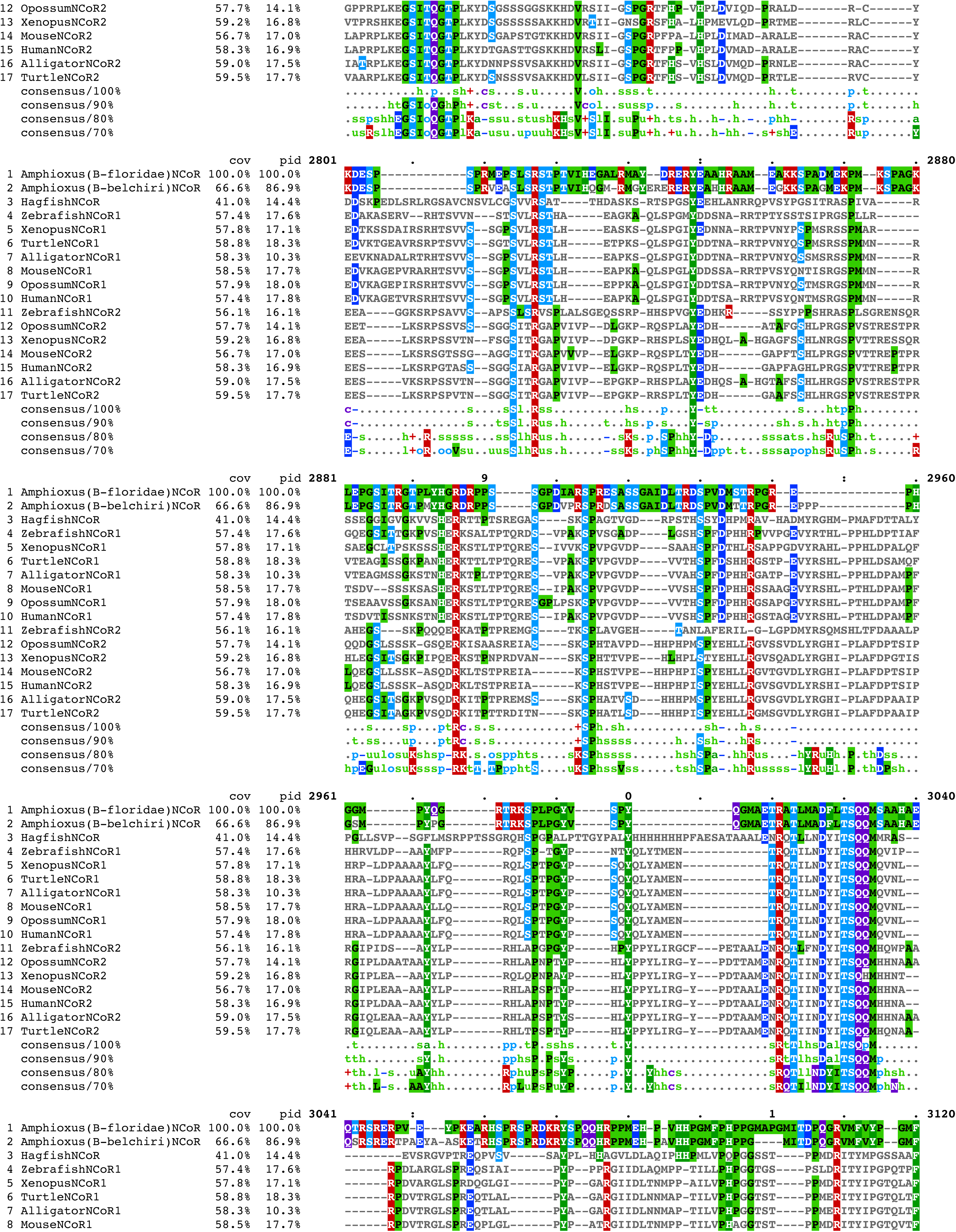
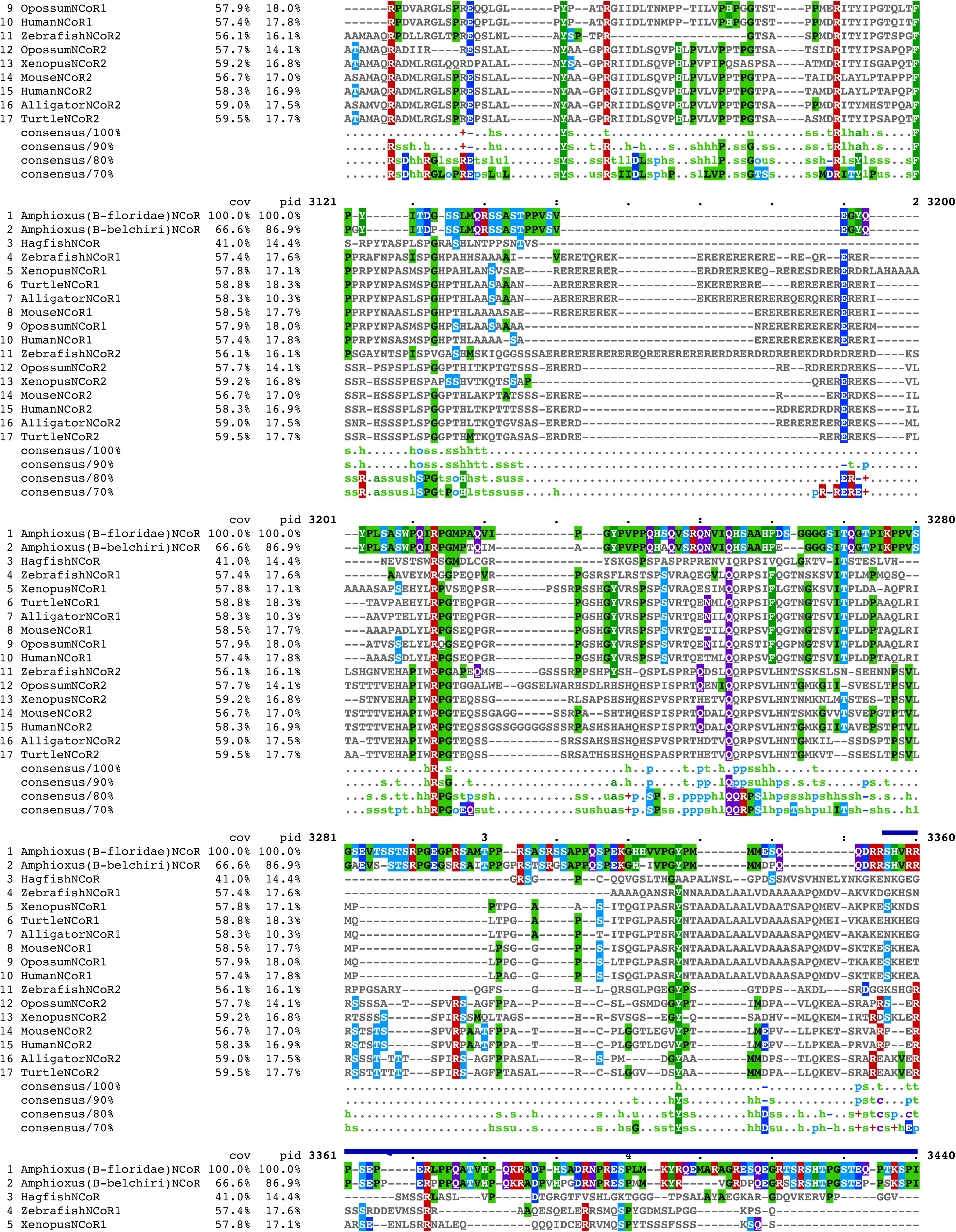
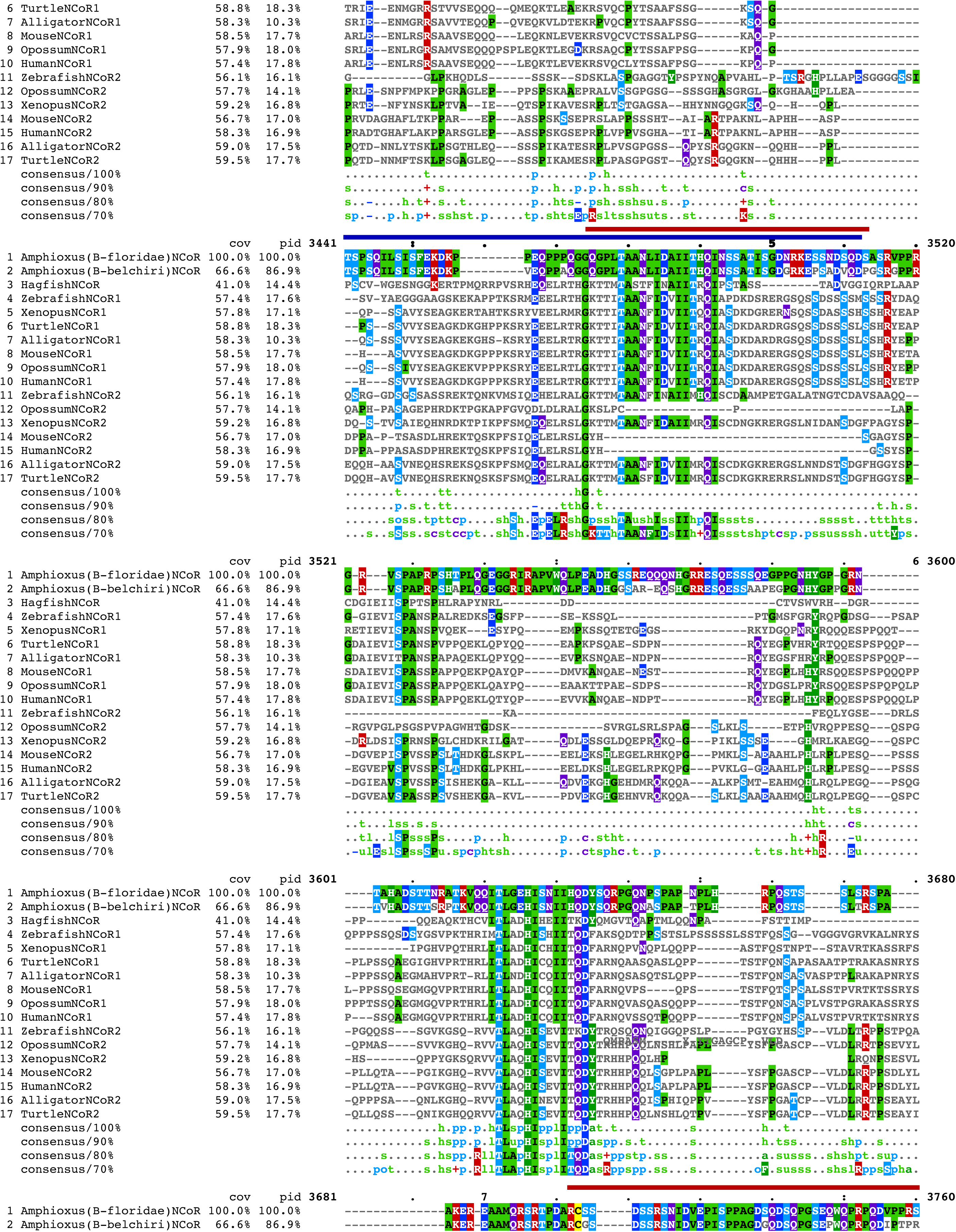
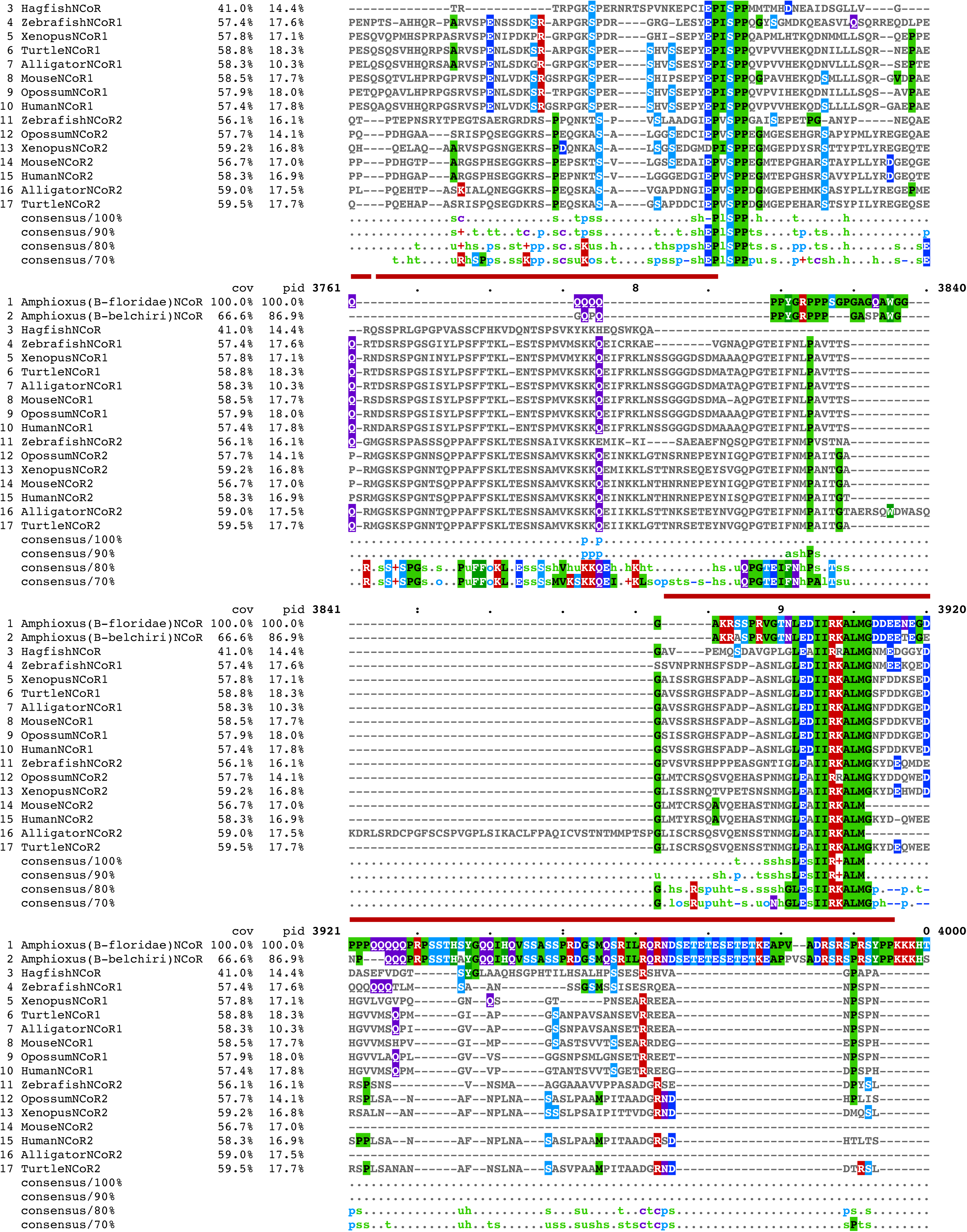
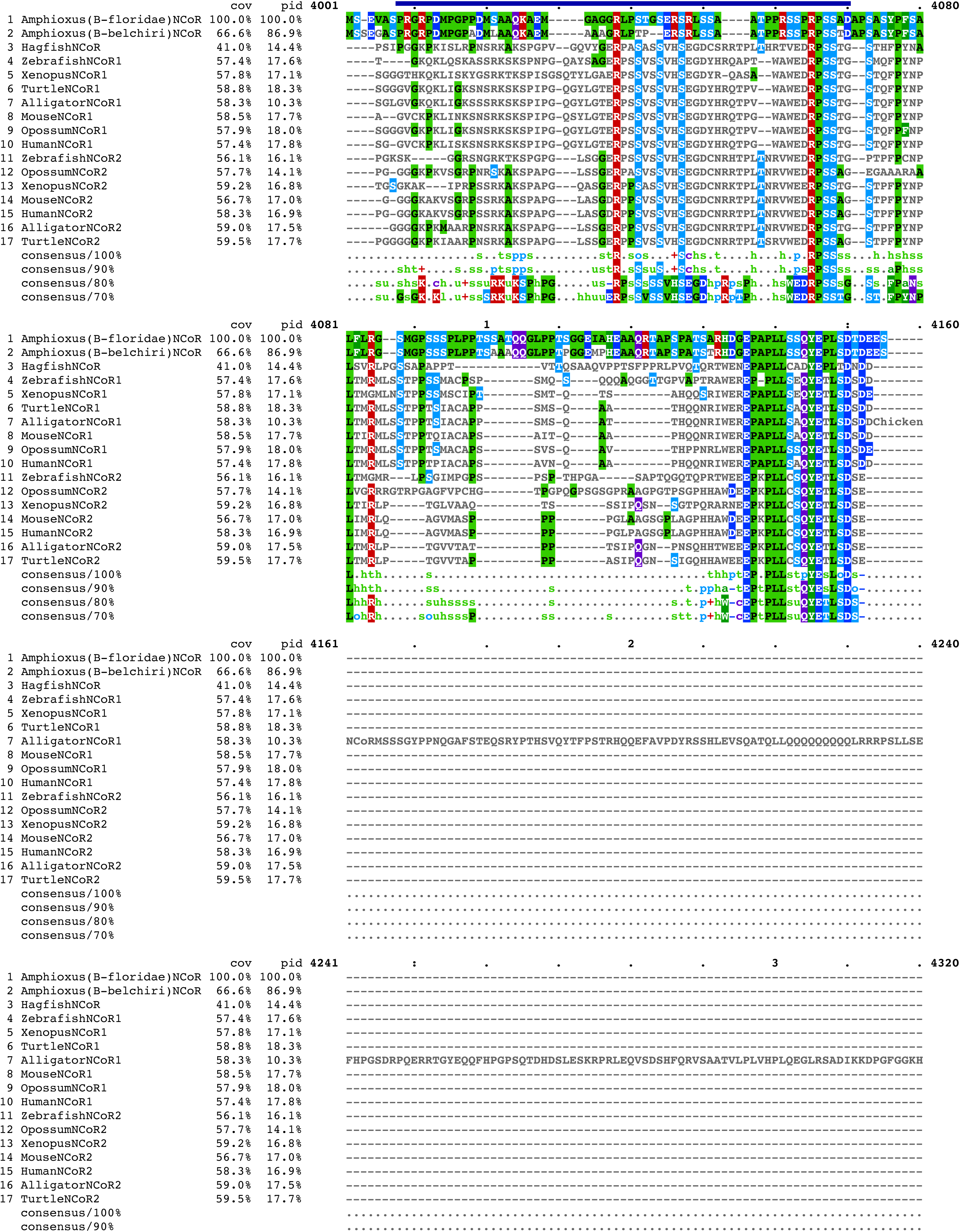
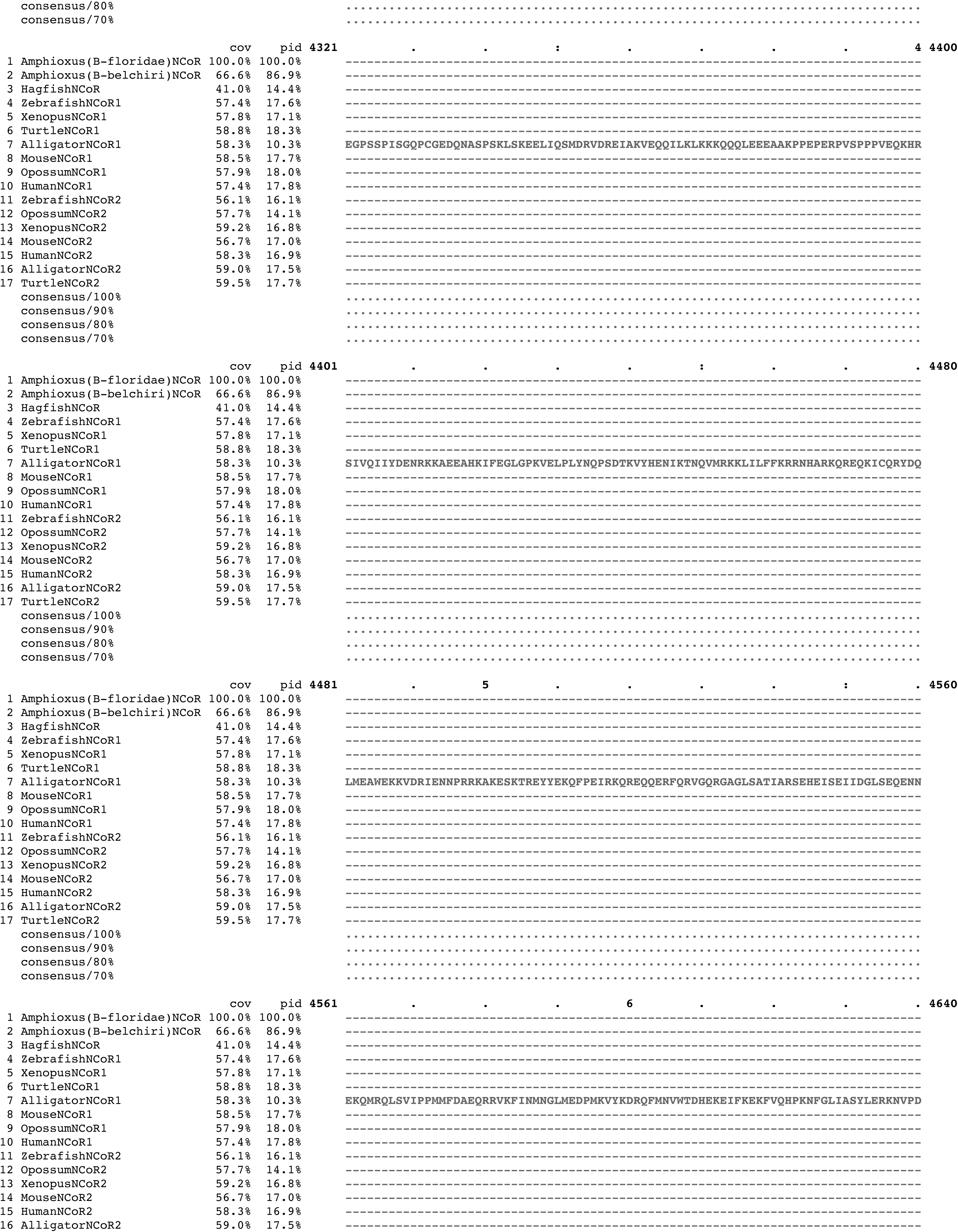
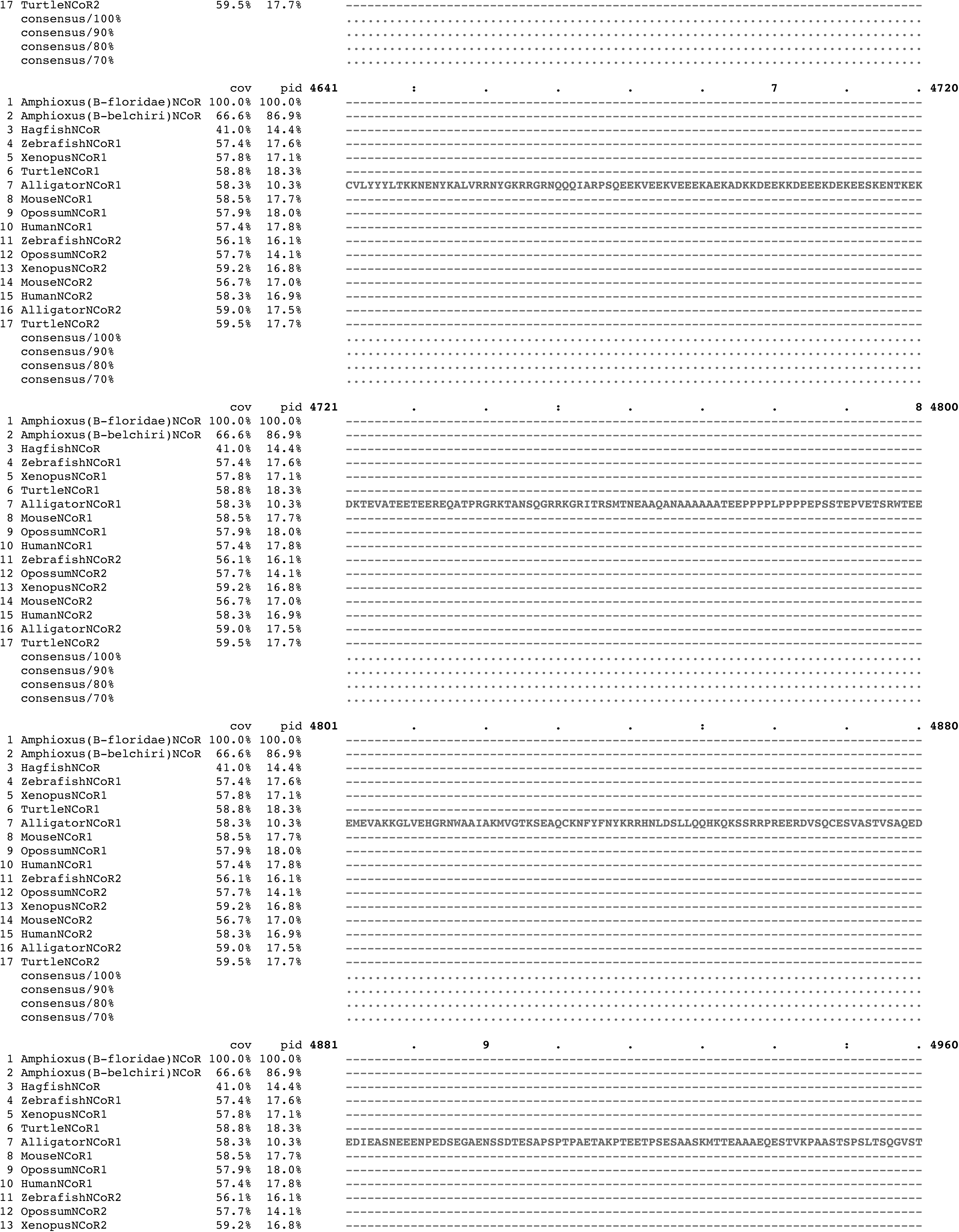
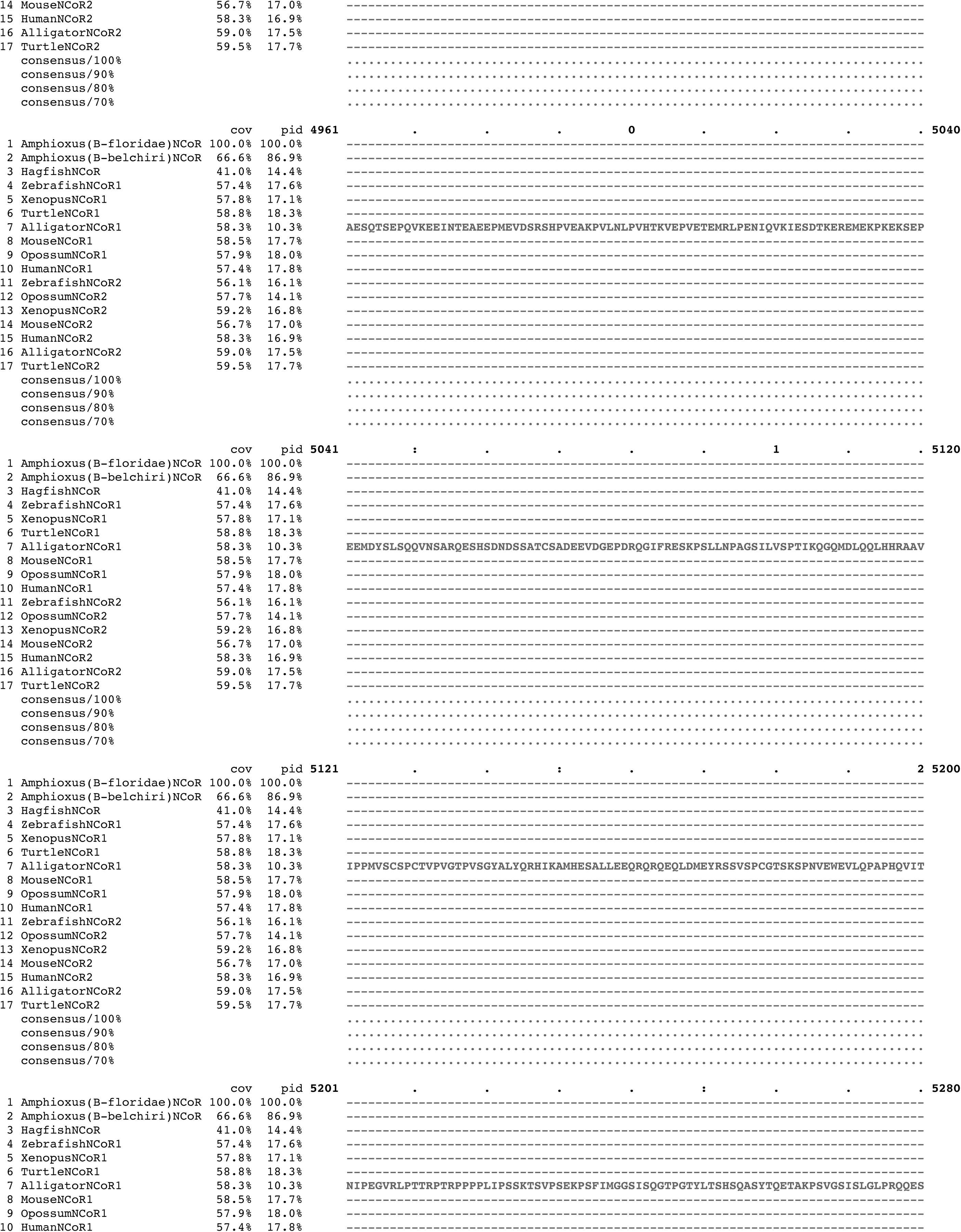
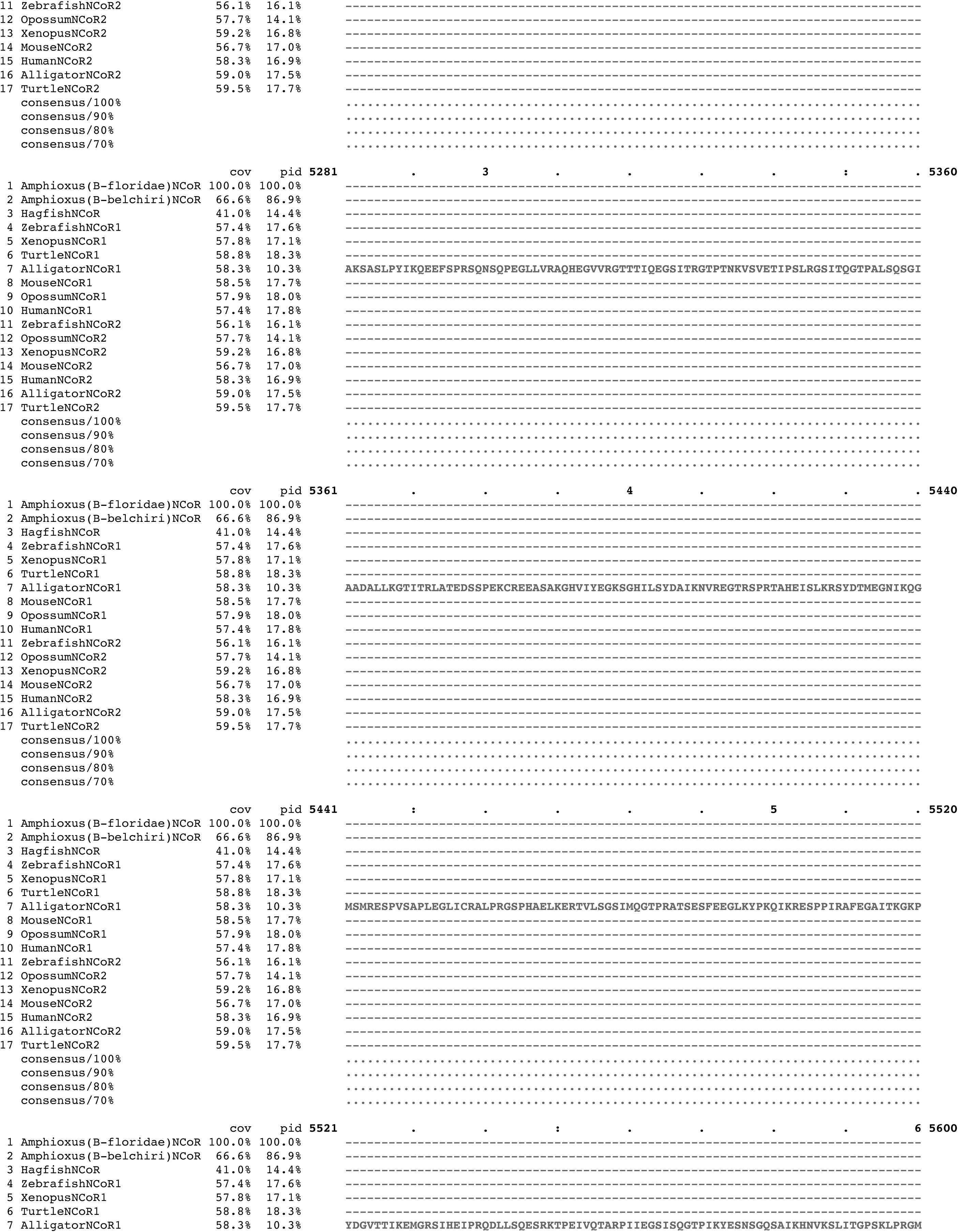
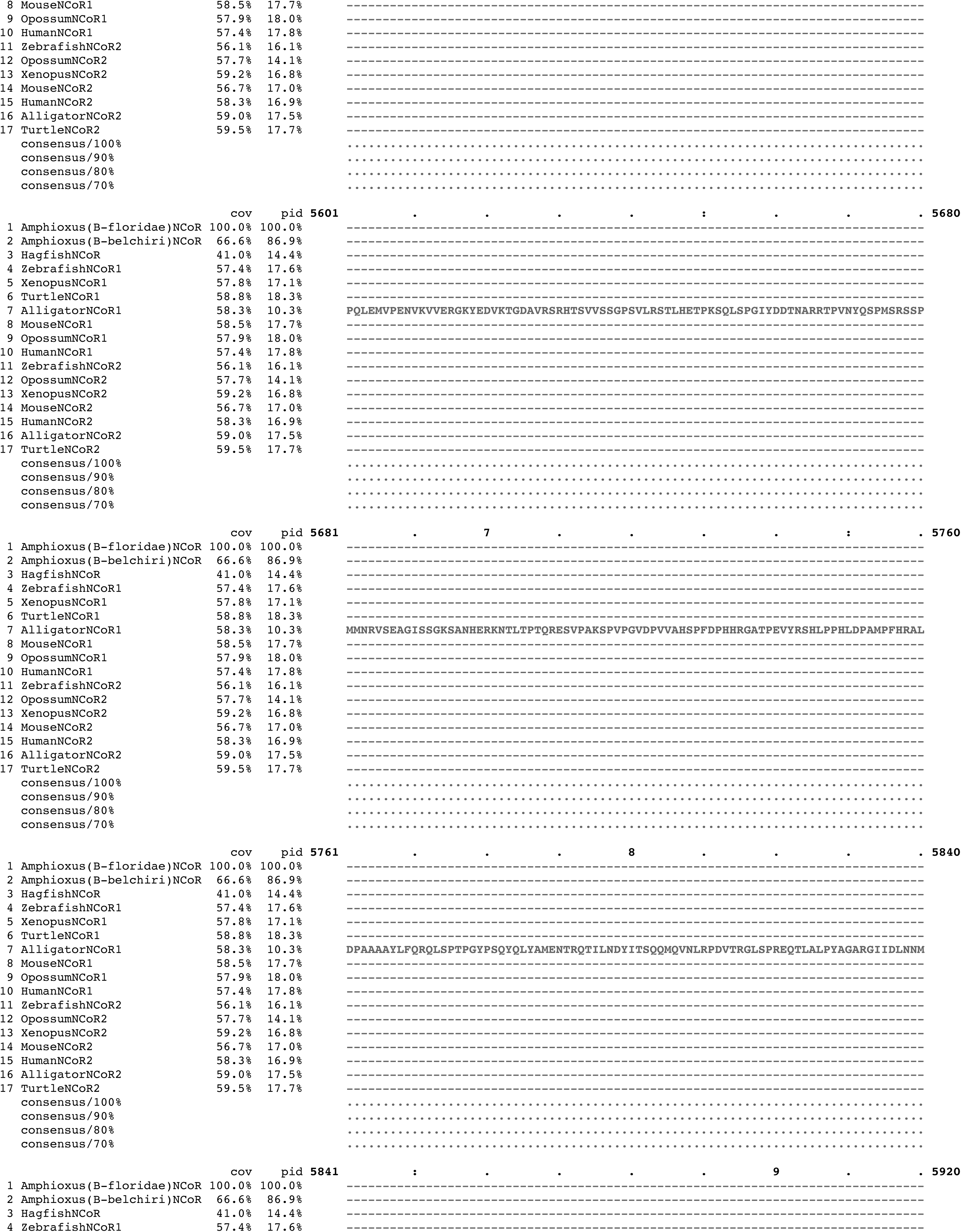
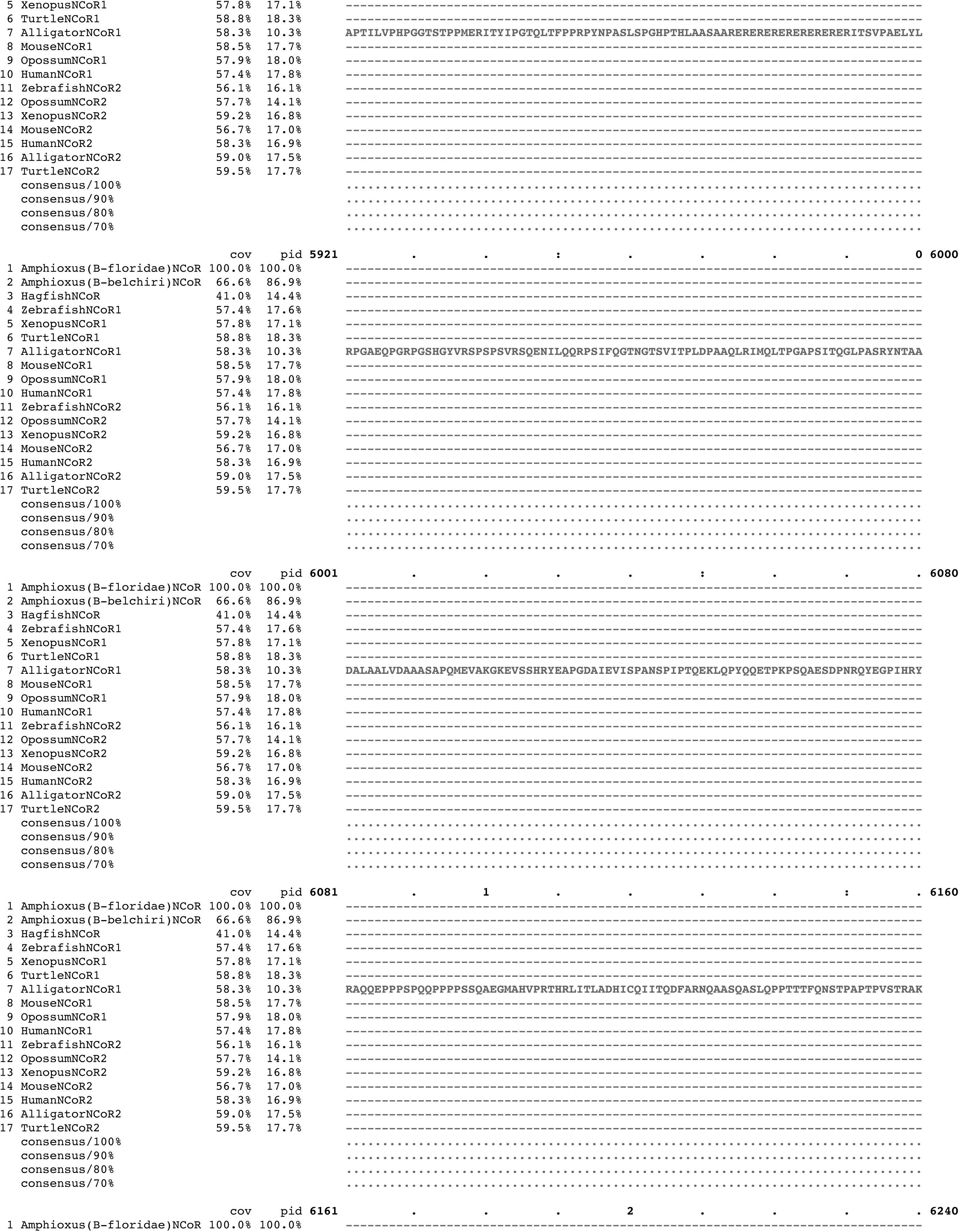
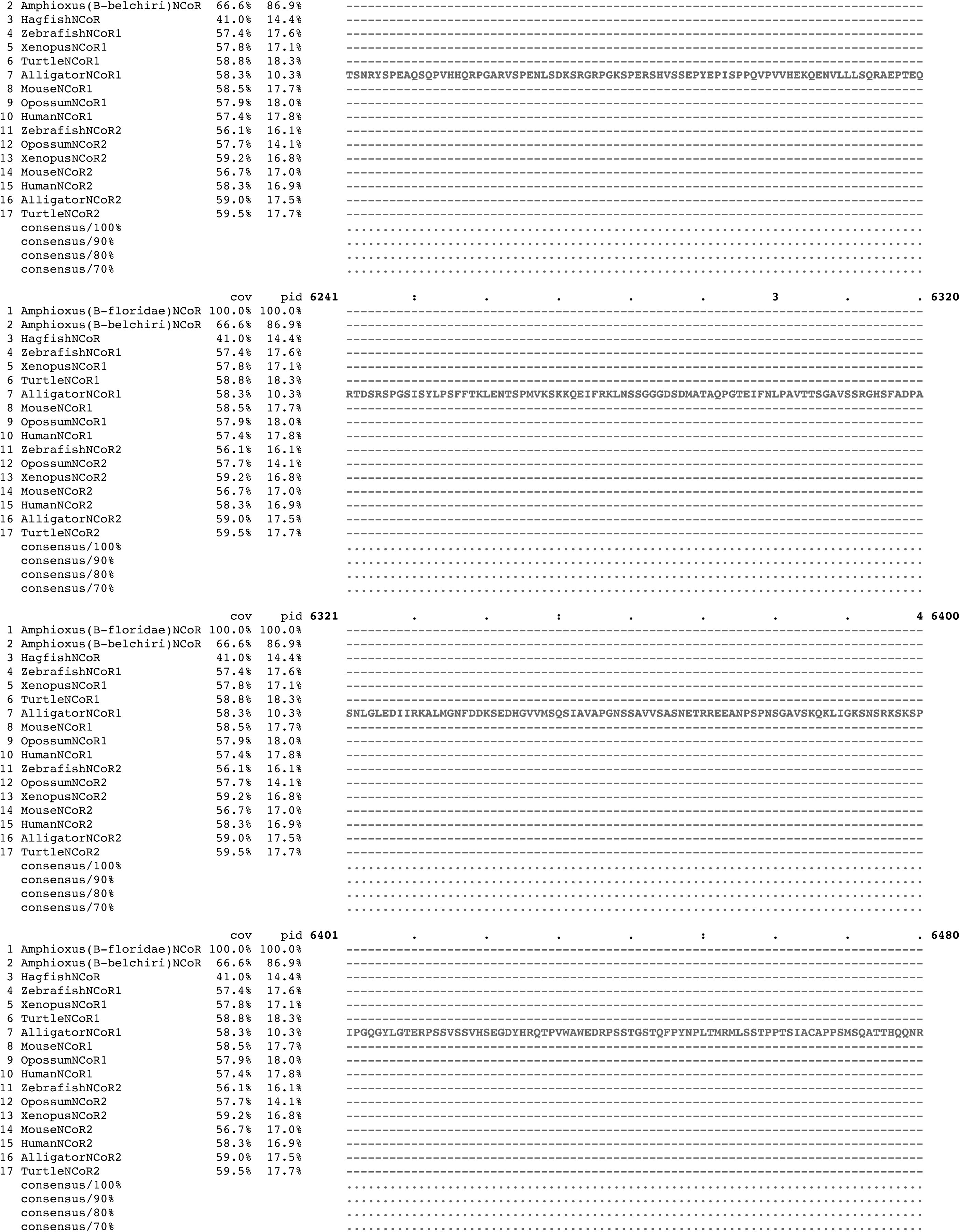
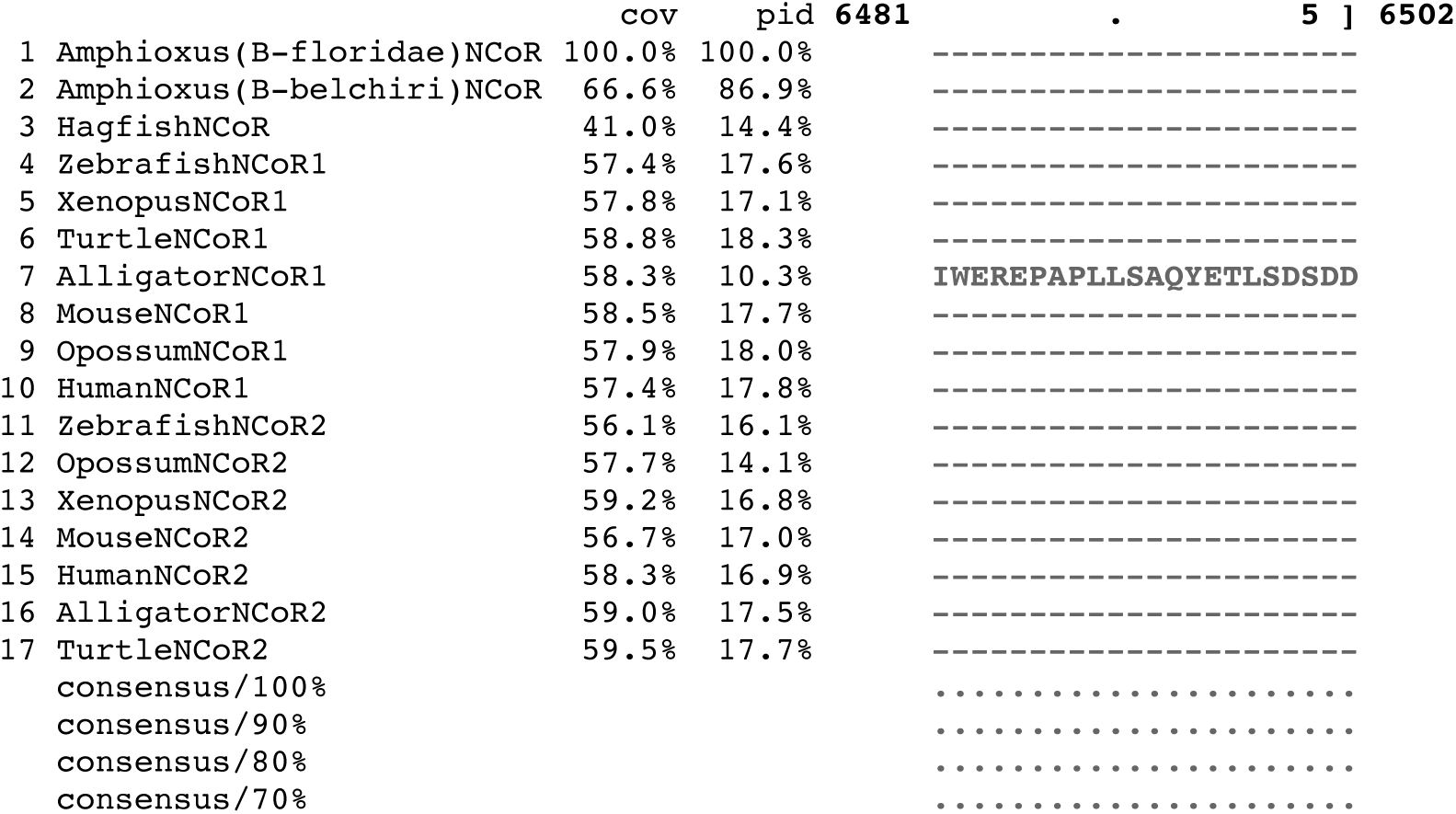
Multispecies alignment of NCoR cDNA sequences. Species are indicated on left. Sequences were aligned using Clustal Omega (https://www.ebi.ac.uk/Tools/msa/clustalo/) and reformatted using Mview (https://www.ebi.ac.uk/Tools/msa/mview/). Color text identifies amino acid class. Dashes are gaps in alignment. Consensus sequences are at bottom. Blue bars at top of sequences represent NCoR-1 regions alternatively spliced in one or more vertebrate; red bars represent NCoR-2 regions alternatively spliced in one or more vertebrate.

